# PKCδ regulates chromatin remodeling and DNA repair through SIRT6

**DOI:** 10.1101/2023.05.24.541991

**Authors:** Trisiani Affandi, Ami Haas, Angela M. Ohm, Gregory M. Wright, Joshua C. Black, Mary E. Reyland

## Abstract

Protein kinase C delta (PKCδ) is a ubiquitous kinase whose function is defined in part by localization to specific cellular compartments. Nuclear PKCδ is both necessary and sufficient for IR-induced apoptosis, while inhibition of PKCδ activity provides radioprotection *in vivo.* How nuclear PKCδ regulates DNA-damage induced cell death is poorly understood. Here we show that PKCδ regulates histone modification, chromatin accessibility, and double stranded break (DSB) repair through a mechanism that requires SIRT6. Overexpression of PKCδ promotes genomic instability and increases DNA damage and apoptosis. Conversely, depletion of PKCδ increases DNA repair via non-homologous end joining (NHEJ) and homologous recombination (HR) as evidenced by more rapid formation of NHEJ (DNA-PK) and HR (Rad51) DNA damage foci, increased expression of repair proteins, and increased repair of NHEJ and HR fluorescent reporter constructs. Nuclease sensitivity indicates that PKCδ depletion is associated with more open chromatin, while overexpression of PKCδ reduces chromatin accessibility. Epiproteome analysis revealed that PKCδ depletion increases chromatin associated H3K36me2, and reduces ribosylation of KDM2A and chromatin bound KDM2A. We identify SIRT6 as a downstream mediator of PKCδ. PKCδ-depleted cells have increased expression of SIRT6, and depletion of SIRT6 reverses the changes in chromatin accessibility, histone modification and NHEJ and HR DNA repair seen with PKCδ-depletion. Furthermore, depletion of SIRT6 reverses radioprotection in PKCδ-depleted cells. Our studies describe a novel pathway whereby PKCδ orchestrates SIRT6- dependent changes in chromatin accessibility to increase DNA repair, and define a mechanism for regulation of radiation-induced apoptosis by PKCδ.

**One Sentence Summary:** Protein kinase C delta modifies chromatin structure via SIRT6 to regulate DNA repair.

## Introduction

Irradiation (IR) is a widely used and highly effective cancer therapy, however, IR damage to tumor-adjacent healthy tissues, particularly in the gut, bone marrow and oral cavity, can result in significant co-morbidities (*1, 2*). For example, the vast majority of patients treated with IR for head and neck cancer will suffer from oral mucositis, and many will have permanent damage to their salivary glands resulting in decreased saliva production, chronic oral infections, and xerostomia (*3–5*). Therapeutic strategies to prevent or mitigate IR damage to tumor-adjacent tissues are very limited, (*6*), hence there is a need to develop therapeutic interventions that can help reduce the side effects from IR without impacting treatment.

Of the DNA lesions induced by IR (*7*), double stranded breaks (DSB) are the most abundant, and can result in loss of genetic information, chromosomal abnormalities, or cell death if left unrepaired (*8, 9*). DSBs are repaired by non-homologous end joining (NHEJ) throughout the cell cycle and by homologous recombination (HR) during S and G2 phases (*10*). Dynamic changes in chromatin structure accompany DNA repair and are mediated by post translational modifications on histones, including acetylation and methylation (*11*). In particular, chromatin relaxation is thought to facilitate DNA repair by increasing accessibility to factors that sense, bind, and repair DSBs (*12*).

Sirtuin 6 (SIRT6), is a chromatin-bound protein from the NAD^+^-dependent deacetylases and ADP-ribosylases family (*13*) which is rapidly recruited to DNA damage sites where it increases chromatin accessibility to enable recruitment of downstream DNA damage response (DDR) factors and efficient DNA repair (*12, 14, 15*). While SIRT6 is widely known for its histone deacetylase activity, it was first identified as a mono-ADP ribosyltransferase enzyme (*16, 17*). Recently, KDM2A has been identified as a substrate of SIRT6 relevant to DNA repair. Upon oxidative stress, SIRT6 stimulates DSB repair by promoting disassociation of KDM2A from chromatin and accumulation of the H3K36me2 mark which is associated with increased DNA repair (*18*).

Our lab has previously shown that protein kinase C delta (PKCδ) is essential for apoptosis in response to DNA-damaging agents, and that depletion or inhibition of PKCδ provides radioprotection *in vivo* (*19–26*). We have shown that nuclear localization of PKCδ is necessary and sufficient for its ability to regulate apoptosis (*21, 22*), implying a critical role for this kinase in the nucleus. Here we have addressed the mechanism(s) that underlies regulation of DNA-damage-induced cell death by PKCδ. Our data shows that PKCδ inhibits DNA repair, and increases DNA damage and genomic instability. Conversely, depletion of PKCδ enhances DNA repair and provides radioprotection through a mechanism that requires SIRT6. Together, our studies define a novel pathway whereby PKCδ, via SIRT6, orchestrates changes in chromatin accessibility to regulate DNA repair. Furthermore, they define a mechanism for regulation of DNA damage-induced cell death by PKCδ, and suggest novel targets for radioprotection.

## RESULTS

### PKCδ induces DNA damage and genomic instability

Genomic instability is a characteristic of most cancers, and can include gene mutations as well as alternations in chromosome structure and number (*27*). We have previously shown that increased expression of PKCδ induces apoptosis, while depletion or inhibition of PKCδ provides radioprotection (*19, 20, 25, 28-31*). To determine if PKCδ promotes DNA damage and genomic instability, we overexpressed PKCδ in ParC5 cells and quantified DNA damage using a neutral comet assay. ParC5 cells were transiently transfected with plasmids that express wild type PKCδ (pPKCδ), the catalytic fragment of PKCδ (pPKCδCF), or nuclear-targeted full length PKCδ (pPKCδNLS) (*21*) (Fig. 1A). The catalytic fragment of PKCδ is generated by caspase cleavage in response to DNA damage and localizes to the nucleus due to exposure of a NLS which is cryptic in the full-length protein (*21, 22*). While caspase cleavage of PKCδ amplifies the apoptotic response, it is not required *per se* (*21*). All three PKCδ constructs increased DNA damage 3-4- fold compared to control cells (Fig. 1B). Similar results were seen using human embryonic kidney (HEK293) cells (Fig. 1C). In this experiment we also transiently expressed a kinase negative mutant of the catalytic fragment of PKCδ (pPKCδCFKN) and a PKCδ catalytic fragment in which the nuclear localization sequence was mutated to exclude it from the nucleus (pPKCδCFNLM). Both mutations completely abolished the ability of the PKCδ catalytic fragment to induce DNA damage (Fig. 1C) indicating that kinase activity and nuclear localization of PKCδ are both required (*21, 31*).

**Fig. 1.**
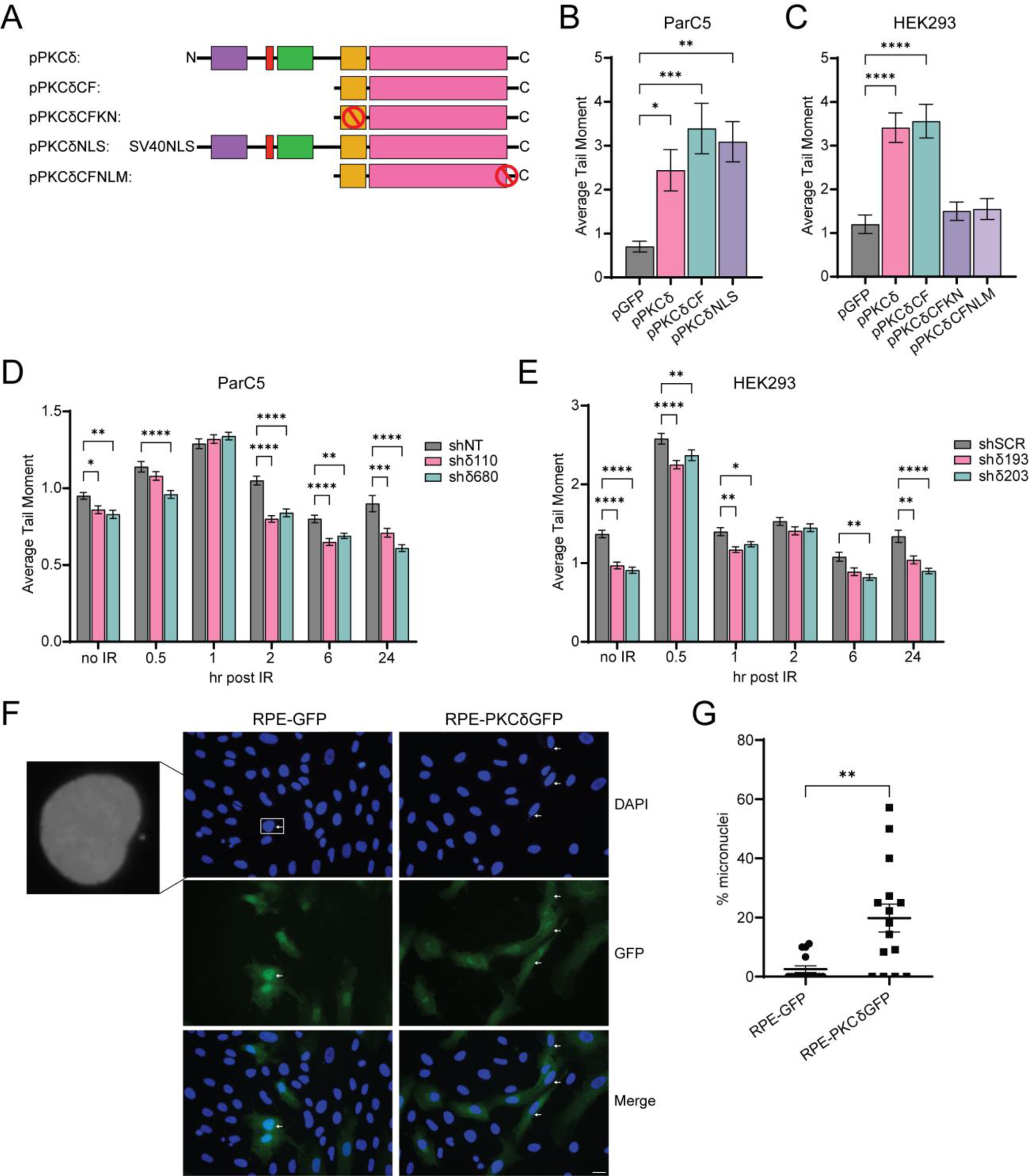
PKCδ leads to genomic instability and induces DNA damage. (**A**) A schematic of the GFP-tagged PKCδ plasmids used, with red circles indicating mutations. (**B**) ParC5 and (**C**) HEK293 cells were transiently transfected with the indicated plasmids for 72 hrs, harvested and assessed for DNA damage using a neutral comet assay. (**D**) ParC5 shNT, shδ110, and shδ680 cells, and (**E**) HEK293 shSCR, shδ193, and shδ203 cells were exposed to 5 Gy IR, harvested at the indicated times and assessed for DNA damage using a neutral comet assay. Data is the mean ± SEM from a representative experiment that was repeated three or more times. (**F**) Representative images of RPE cells stably expressing pGFP or pPKCδGFP. The white arrows indicate micronuclei. Scale bar = 20 µm. (**G**) The percent micronuclei was determined by normalizing the number of micronuclei (*N* = 20 fields/cell line) to the total number of nuclei identified by DAPI staining. Statistics represent one-way ANOVA in (B) and (C) followed by Dunnett’s multiple comparisons to the pGFP controls, two-way ANOVA in (D) and (E) followed by Dunnett’s multiple comparisons within each time point to the corresponding shNT or shSCR control and unpaired t-test in (G). \**P* < 0.05, \*\**P* < 0.01, \*\*\**P* < 0.001, and \*\*\*\**P* < 0.0001.

As depletion of PKCδ suppresses DNA damage induced apoptosis, we asked if IR-induced DNA damage resolves faster in PKCδ-depleted cells. Surprisingly, the amount of DNA damage in non-irradiated PKCδ-depleted cells (shδ110 and shδ680) was significantly less than that observed in ParC5 shNT cells (Fig. 1D, no IR). A similar trend was seen in PKCδ-depleted HEK293 cells, in which two PKCδ-targeting shRNAs, shδ193 and shδ203, had significantly less DNA damage than the shSCR control under basal conditions (Fig 1E, no IR). Following IR, the kinetics of induction and resolution of DNA damage was similar in ParC5 shNT and PKCδ-depleted shδ110 and shδ680 cells, however there was more complete resolution of IR-induced DNA damage in the two PKCδ-depleted cell lines compared to control shNT cells (Fig. 1D). PKCδ-depleted HEK293 cells (shδ193 and shδ203), also show enhanced resolution of IR-induced DNA damage as compared to the control shSCR cells (Fig. 1E). These data indicate that radioprotection mediated by depletion of PKCδ is associated with more complete resolution of DNA damage.

Genomic instability can arise due to impaired DNA DSB repair (*32*). As micronuclei are a biomarker for DNA damage and chromosomal instability (*33*), we quantified micronuclei formation in human retinal pigment epithelial (RPE) cells stably integrated with pGFP or pGFP- tagged PKCδ (RPE-GFP or RPE-PKCδGFP cell lines, respectively). We observed a 10-fold increase in micronuclei formation in RPE-PKCδGFP cells compared to control RPE-GFP cells (Fig. 1, F and G), indicating that increased expression of PKCδ is sufficient to cause genomic instability in immortalized, but non-transformed normal epithelial cells. We next interrogated the TCGA database to determine if differences in expression of PKCδ correlate with copy number variants (CNV) in cancer patients with Lung Adenocarcinoma (LUAD) or Head-Neck Squamous Cell Carcinoma (HNSCC). Surprisingly, high PKCδ expression correlated with low CNV in both LUAD and HNSCC cancers (Fig. S1, A and B). Thus, it is likely that in these tumors, increased DNA instability is coupled with increased apoptosis, suggesting a tumor suppressive role for PKCδ.

### PKCδ regulates NHEJ and HR mediated DNA DSB repair

To determine if PKCδ regulates NHEJ and HR repair pathways we generated ParC5 reporter cell lines (shNT, shδ110 and shδ680) and HEK293 reporter cell lines (shSCR, shδ193, shδ203 and shδ800) containing either chromosomally integrated GFP-based NHEJ or HR reporter constructs (*34, 35*). In this assay, successful NHEJ or HR repair results in GFP fluorescence. Each reporter line was co-transfected with pI-SceI to induce DSB and pDsRed to normalize for the differences in transfection efficiency between cell lines. All PKCδ-depleted cell lines showed an increase in both NHEJ and HR mediated DNA repair (Fig. 2, A to C). In ParC5 NHEJ reporter cells, repair increased 3.5- and 1.5-fold in shδ110 and shδ680, respectively, compared to the shNT control (Fig. 2A). Similarly, in two PKCδ-depleted HEK293 NHEJ reporter cell lines (shδ193, shδ800) (Fig. S2A), repair increased over 3-fold compared to the shSCR control (Fig. 2B). Depletion of PKCδ also increased HR repair in all four PKCδ-depleted cell lines, although in general the increase in HR mediated repair was of a lower magnitude than the increase in NHEJ mediated DNA repair for both ParC5 and HEK293 cells (Fig. 2, A and C).

**Fig. 2.**
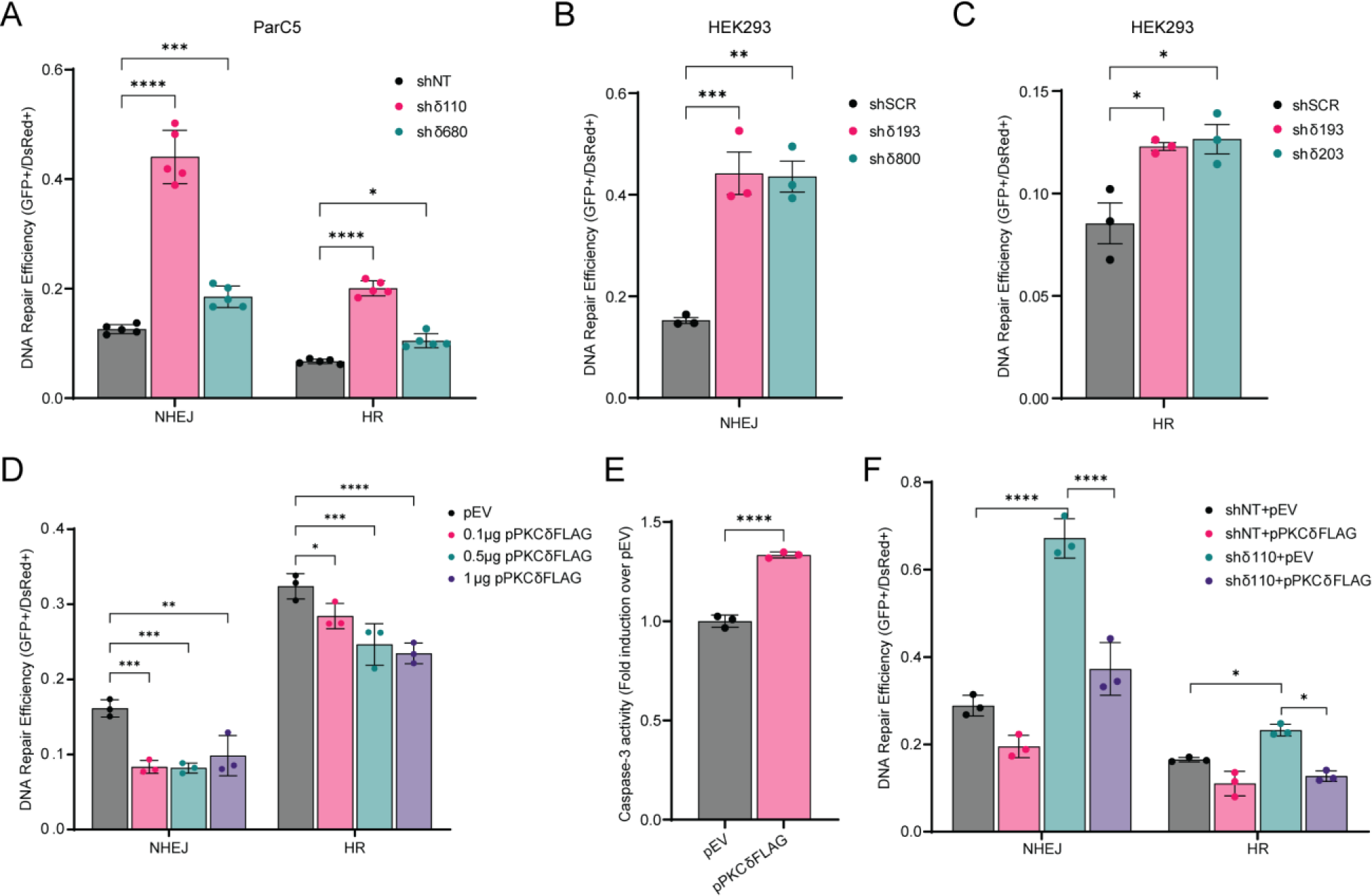
PKCδ regulates NHEJ and HR mediated DSB repair. (**A to C**) DNA repair was analyzed in (A) ParC5 shNT, shδ110, and shδ680 cells containing stably integrated NHEJ or HR reporters, (B) HEK293 shSCR, shδ193, and shδ800 cells containing NHEJ reporters, or (C) HEK293 shSCR, shδ193, and shδ203 cells containing HR reporters. Cells were co-transfected with plasmids expressing pI-SceI to induce DSBs and pDsRed to normalize transfection efficiency. Reporter cells were allowed to grow for 72 hrs post-electroporation and repair was quantified by flow cytometry. (**D**) ParC5 NHEJ or HR reporter cells were transfected with 1 µg pEV (black bars) or increasing amount of pPKCδFLAG, and DNA repair was assayed. (**E**) ParC5 cells transfected with 0.1 ug of PKCδFLAG for 72 hrs were collected, lysed, and assayed for caspase-3 activities. (**F**) ParC5 shNT and shδ110 reporter cells were co-transfected with plasmids expressing pI-SceI, pDsRed, and pEV or pPKCδFLAG and DNA repair was analyzed. Each data point represents an average from three independent biological replicates. Data is shown as mean ± SEM. Statistics represent two-way ANOVA or one-way ANOVA for (B) and (C), followed by Dunnett’s multiple comparisons within NHEJ or HR reporter cell lines to their corresponding shNT, shSCR, or pEV control, and unpaired t-test for (E). \**P* < 0.05, \*\**P* < 0.01, \*\*\**P* < 0.001, and \*\*\*\**P* < 0.0001.

We show that in the absence of PKCδ, NHEJ and HR mediated DNA repair is increased, indicating that PKCδ is a negative regulator of DSB repair. To explore this further, we expressed increasing amounts of FLAG-tagged PKCδ (pPKCδFLAG) in ParC5-NHEJ and ParC5-HR reporter cells. Expression of pPKCδFLAG decreased both NHEJ and HR mediated DNA repair, albeit NHEJ reporter cells appeared to be more sensitive (Fig. 2D and Fig. S2B). Decreased repair correlates with an increase in apoptosis in NHEJ reporter cells (Fig. 2E). Similarly, transfection of pPKCδFLAG reversed the increase in NHEJ and HR repair seen in shδ110 reporter cells (Fig. 2F and Fig. S2C). Together our data indicates that PKCδ regulates DNA damage-induced apoptosis at least in part through regulation of DNA repair.

### Loss of PKCδ “primes” cells for DSB repair

A crucial mechanism for cells to maintain genomic integrity in the face of endogenous insults and exogenous damaging agents is through activation of DDR, which senses DNA damage and promotes repair (*32*). One of the earliest events in the cellular response to DSBs is the phosphorylation of the histone protein H2AX at Ser^139^ (γH2AX), which accumulates at sites of DSBs to recruit and localize DNA repair proteins (*36*). To examine the kinetics of γH2AX foci formation, foci were quantified in ParC5 shNT, shδ110 and shδ680 cells following exposure to IR. γH2AX foci increased in all cell lines at 0.5 hr, however PKCδ-depleted shδ110 and shδ680 cells had significantly fewer foci compared to shNT cells (Fig. 3A). Moreover, γH2AX foci resolved sooner in shδ110 and shδ680 cells (Fig. 3A, 6 hr and 24 hr), indicating faster DNA repair. To investigate the kinetics of NHEJ and HR repair directly, we assayed DNA repair foci using antibodies to specific repair complex proteins. Surprisingly, in PKCδ-depleted cells, both NHEJ (DNA-PK pSer^2056^) and HR (Rad51) foci (Fig. 3, B and C) are increased basally, which correlates with the reduction in DNA damage previously observed (Fig. 1D). Following IR, PKCδ-depleted cells have more DNA-PK pSer^2056^ foci at 0.5 hr, but these resolve faster than in shNT cells (Fig. 3B, 0.5 hr versus 4 hr). Similarly, there were more Rad51 foci at 2 hr in PKCδ-depleted cells compared to shNT cells (Fig. 3C). These data suggest that in the absence of PKCδ, cells are primed to repair DSBs.

**Fig. 3.**
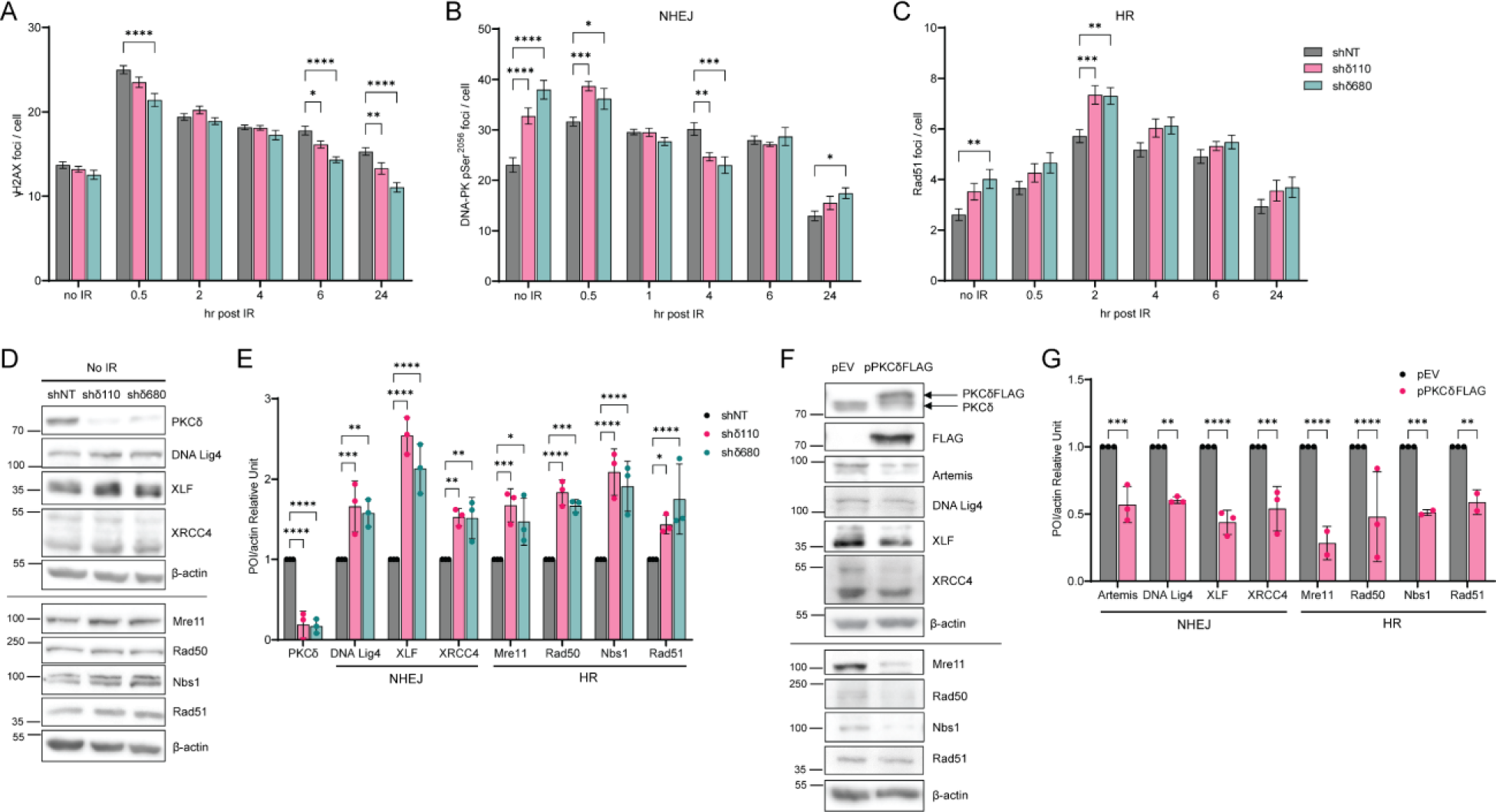
Loss of PKCδ enhances formation of DNA repair foci and alters expression of NHEJ and HR repair proteins. (**A to C**) ParC5 shNT, shδ110, and shδ680 cells were plated on coverslips and exposed to 1 Gy IR. Cells were harvested at indicated times post IR by permeabilization, fixation, and stained for foci, (A) γH2AX, (B) DNA-PK pSer^2056^, and (C) Rad51 foci (legend in C is for A to C). An average of three independent experiments is shown. Error bars show ± SEM. (**D**) Western blots of ParC5 shNT, shδ110, and shδ680 cells immunoblotted with the indicated antibodies. (**E**) Densitometry of Western blots shown in D; expression of each protein of interest (POI) was normalized to the actin control. (**F**) ParC5 cells were transiently transfected with empty vector (pEV) or pPKCδFLAG for 48 hrs. Cells were then collected and immunoblotted with the indicated antibodies (First and second blots were probed with antibodies against PKCδ and FLAG-tag respectively). Note that panels D and F show two different experiments separated by lines, each normalized to their own β-actin loading control. (**G**) Densitometry of Western blots shown in F; expression of each POI was normalized to the actin control. Data are from three independent biological replicates and shown as mean ± SEM. Statistics represent two-way ANOVA with Dunnett’s multiple comparisons and Sidak’s multiple comparisons for (G) to the corresponding control (black bars). \**P* < 0.05, \*\**P* < 0.01, \*\*\**P* < 0.001, and \*\*\*\**P* < 0.0001.

To investigate the mechanism(s) underlying priming of DSB repair, we looked at expression of a subset of NHEJ and HR repair proteins. All NHEJ and HR repair proteins assayed increased in shδ110 and shδ680 cells compared to shNT cells, ranging from a 1.4-fold (Rad51) to a 2.5-fold increase (DNA Lig4) (Fig. 3, D and E). Similar results were seen when mRNA abundance was quantified by qRT-PCR (Fig. S3A). On the contrary, when pPKCδFLAG was overexpressed in ParC5 cells, we observed a dramatic decrease in expression of NHEJ and HR repair proteins (Fig. 3, F and G). The mRNA levels of these proteins were similarly decreased (Fig. S3B). Thus, changes in the expression of NHEJ and HR repair proteins likely contribute to priming of DSB repair pathways by PKCδ.

### PKCδ regulates global chromatin structure

Both NHEJ and HR repair require a dynamic switch from condensed to relaxed chromatin to provide accessibility of DNA repair proteins to the site of DSBs. To determine if PKCδ affects global chromatin structure, we utilized a micrococcal nuclease (MNase)-digested chromatin assay (*37–39*). PKCδ-depleted ParC5 and HEK293 cells showed increased elution of DNA at a lower molarity of NaCl compared to their respective control shRNA cell lines (Fig. 4, A and B). The relative percent of DNA eluted at 80 mM NaCl increased by 1.2 and 1.3-fold in shδ110 and shδ680 cells, respectively, compared to shNT control cells (Fig. 4A), suggesting more accessible chromatin in the absence of PKCδ. Similar results were seen in PKCδ-depleted HEK293 cells (Fig. 4B). To verify that PKCδ kinase activity and nuclear localization are required for changes in chromatin structure, ParC5 shδ110 cells were transiently transfected with pPKCδCF or pPKCδCFKN (Fig. 4C), or pPKCδNLS or pPKCδNLM (Fig. 4D) and MNase sensitivity was compared to shNT or shδ110 cells transfected with pGFP alone. As expected, in shδ110+pGFP transfected cells more DNA was eluted in the 80 mM NaCl fraction compared to the shNT+pGFP (Fig. 4, C and D), consistent with what we observed in Fig. 4A and 4B. However, expression of pPKCδCF in shδ110 cells significantly reduced the percent of DNA eluted in the 80 mM NaCl fraction when compared to the shδ110+GFP (Fig. 4C), while no change in MNase sensitivity was seen with expression of the catalytically negative pPKCδCFKN (Fig. 4C), verifying that the kinase activity of PKCδ is required to induce chromatin condensation. Similarly, expression of pPKCδNLS, but not pPKCδNLM, reduced elution of DNA at 80 mM compared to shδ110+pGFP transfected cells (Fig. 4D), verifying that the nuclear localization of PKCδ is required to induce chromatin condensation. These data implicate PKCδ as a regulator of global chromatin structure, consistent with its ability to regulate DNA repair pathways. Furthermore, they suggest that depletion of PKCδ likely primes cells for DNA repair by increasing the accessibility of repair proteins to sites of DSBs.

**Fig. 4.**
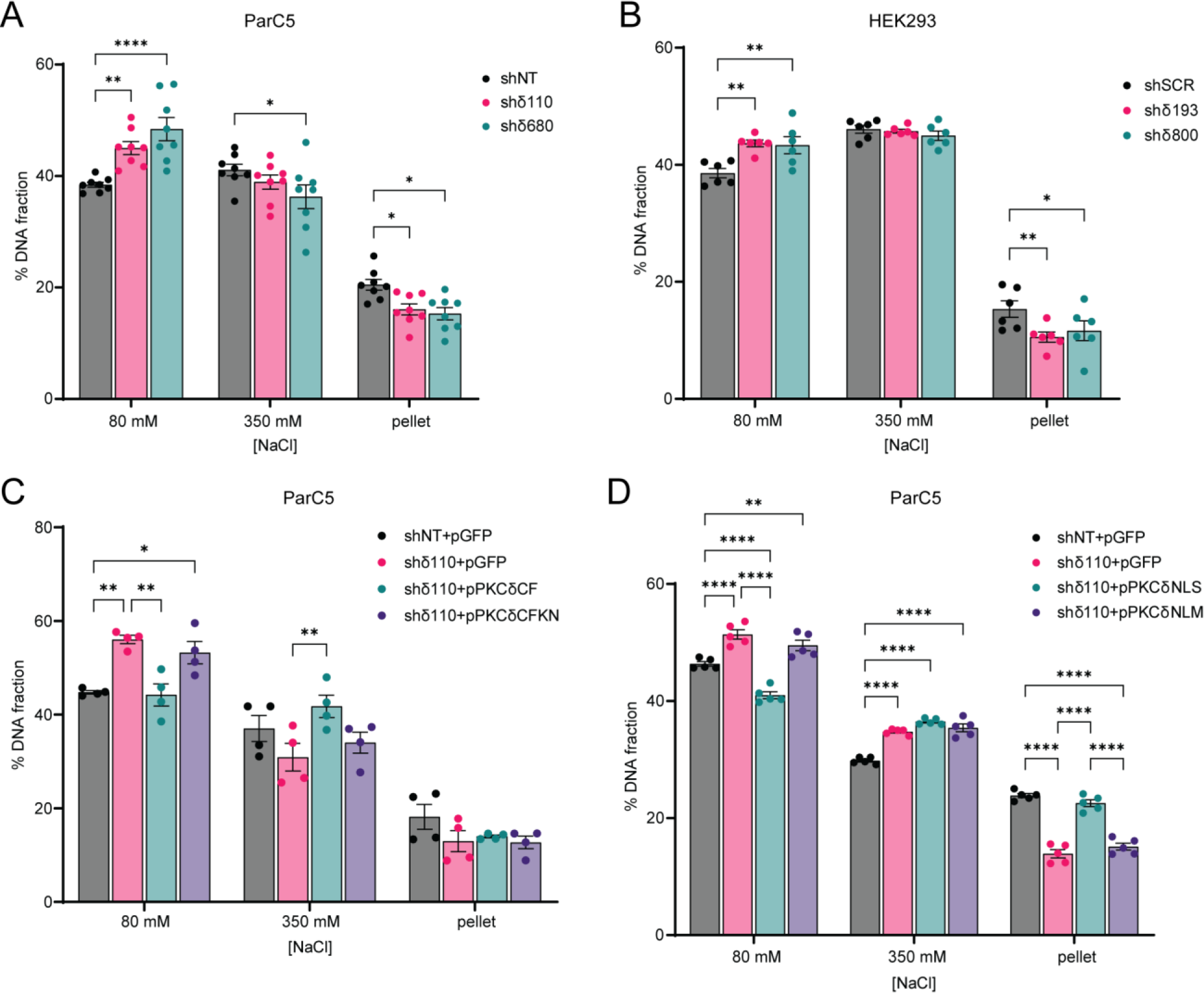
PKCδ alters global chromatin structure. (**A and B**) MNase assay was performed on (A) ParC5 shNT, shδ110, and shδ680 cells, and on (B) HEK293 shSCR, shδ193, and shδ800 cells. (**C and D**) ParC5 shNT cells were transfected with pGFP only, while shδ110 cells were transfected with (C) pGFP, pPKCδCF, or pPKCδCFKN, or (D) pGFP, pPKCδNLS, or pPKCδNLM for 24 hrs prior to harvesting. Percentage of DNA in each [NaCl] fraction was assayed as described in Materials and Methods. Shown is data (mean ± SEM) from at least three independent biological replicates. Statistics represent two-way ANOVA followed by Dunnett’s multiple comparisons within each fraction to their corresponding shNT or shSCR control for (A) and (B), or two-way ANOVA followed by Tukey’s multiple comparisons within each fraction for (C) and (D). \**P* < 0.05, \*\**P* < 0.01, \*\*\**P* < 0.001, and \*\*\*\**P* < 0.0001.

### Depletion of PKCδ is associated with increased H3K36 methylation

Methylation and acetylation of histones can regulate the efficiency of DNA repair by controlling the accessibility of damaged DNA to repair factors. To investigate the contribution of chromatin regulating enzymes and histone PTMs to changes in chromatin accessibility in PKCδ- depleted cells, we first used general inhibitors that target a subset of these enzymes. These include Trichostatin A (TSA) to inhibit Class I and II histone deacetylases (HDACs), nicotinamide (NAM) to inhibit Class III HDACs (SIRT1-7), and succinate (SUC) to inhibit jumonji demethylases (KDMs). No changes in NHEJ repair efficiency were observed when shNT cells were treated with either TSA or NAM (Fig. 5A). However, treatment of shδ110 cells with TSA or NAM resulted in a 50% decrease in NHEJ repair compared to the untreated (UT) shδ110 cells (Fig. 5A), implying that Class I, II or III HDACs are involved in promoting DNA repair in PKCδ-depleted cells. Similarly, SUC had no effect on NHEJ repair in shNT cells, but further increased repair in shδ110 cells compared to UT cells (Fig. 5B), implicating KDMs as inhibitors of NHEJ in shδ110 cells. This suggests that increased repair in shδ110 cells may require decreased histone acetylation and increased methylation. We next looked at the effect of these inhibitors on chromatin structure using the MNase assay. NAM reversed the chromatin relaxation seen in shδ110 cells, as evidenced by a reduction in DNA elution at 80 mM NaCl and an increase in DNA elution at 350 mM NaCl (Fig. 5C). However, SUC did not increase DNA elution at 80 mM NaCl beyond that seen in shδ110 cells (Fig. 5D).

**Fig. 5.**
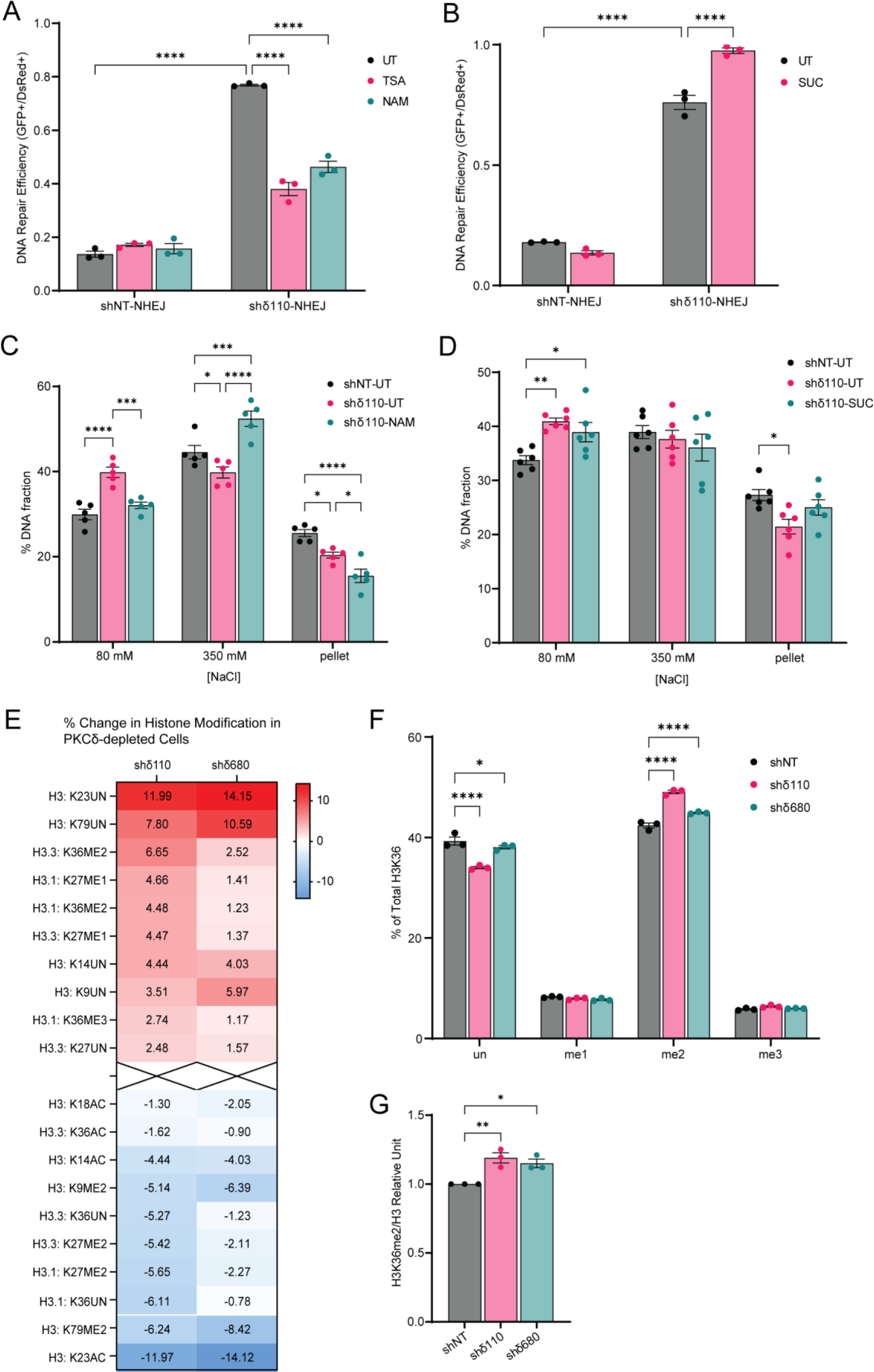
Depletion of PKCδ promotes DNA repair and chromatin relaxation and increases H3K36me2. (**A and B**) DNA repair was analyzed in ParC5 shNT-NHEJ and shδ110-NHEJ reporter cells pretreated with (A) 100 nM trichostatin A (TSA) or 5 mM nicotinamide (NAM), or (B) 5 mM succinate (SUC) for 24 hrs and then co-transfected with plasmids expressing pI-SceI to induce DSBs and pDsRed to normalize transfection efficiency. NHEJ and HR reporter cells were allowed to grow for 72 hrs post-transfection and repair was quantified. (**C and D**) MNase assay was performed on ParC5 shNT and shδ110 cells pretreated with (C) 5 mM NAM or (D) 5 mM SUC for 48 hrs before harvesting. Percentage of DNA in each [NaCl] fraction was assayed. Shown is data (mean ± SEM) from at least three independent biological replicates. (**E**) Histone modifications of ParC5 shNT, shδ110, and shδ680 were analyzed by mass spectrometry (Northwestern University). Heat map shows the percent change of PTMs on H3 in each modification in shδ110 and shδ680 cells relative to shNT (top and bottom 10 shown). (**F**) Profile of H3K36 methylation in shNT, shδ110, and shδ680 cells. (**G**) Densitometry of H3K36me2 normalized to total H3. Shown is data (mean ± SEM) from three independent biological replicates. Statistics represent two-way ANOVA followed by Sidak’s for (A), Tukey’s for (B to D), and Dunnett’s multiple comparisons for (F), and one-way ANOVA followed by Dunnett’s multiple comparisons for (G). \**P* < 0.05, \*\**P* < 0.01, \*\*\**P* < 0.001, and \*\*\*\**P* < 0.0001.

As PKCδ regulation of chromatin structure and DNA repair could be mediated by epigenetic modifications, we profiled histone modifications in ParC5 shNT and PKCδ-depleted shδ110 and shδ680 cells using the Epiproteomic Histone Modification Panel (EHMP) in collaboration with the Northwestern Proteomics Core Facility. Acid-extracted histones were identified by mass spectrometry, and the percentage change in each histone PTM between the shδ110, shδ680 cells and shNT cells were calculated and plotted in the heatmap (Fig. 5E, top and bottom 10 of PTMs on H3 are shown). Full EHMP results are available in Supplementary Materials (Data file S1). As predicted by our inhibitor studies (Fig. 5, A to D), depletion of PKCδ correlated with a reduction in histone acetylation (Fig. 5E, H3K23, H3K14, H3K36 and H3K18). Changes in histone methylation were more varied and included a decrease in H3K79me2, H3K27me2, and H3K9me2, and an increase in H3K27me1 (Fig. 5E). However, the largest increase in histone methylation observed was at H3K36me2 (Fig. 5, E and F). Of the histone marks modified in a PKCδ-dependent manner, H3K36me2, H3K79me2 and H3K14ac are associated with increased DNA repair (*40–43*). As H3K79me2 and H3K14ac are reduced in PKCδ-depleted cells, they are unlikely to be involved in the phenotypes we have described. Therefore, we focused on the H3K36me2 mark as a mechanism to explain increased DNA repair in PKCδ-depleted cells. We confirmed enrichment of H3K36me2 in PKCδ-depleted cells by immunoblot of acid-extracted histones with an antibody specific to dimethyl H3K36. In agreement with the EHMP results, H3K36me2 were increased up to 15% in shδ110 and shδ680 cells (Fig. 5G).

### PKCδ regulates chromatin, DSB repair and apoptosis through SIRT6

SIRT6 is an early DSB sensor that facilitates opening of chromatin to enable recruitment of DNA repair factors to sites of damage (*13*). Rezazadeh *et al.* have shown that SIRT6 can also increase H3K36me2 at DNA damage sites by displacement of KDM2A from chromatin through a mechanism dependent on ribosylation of KDM2A by SIRT6 (*18*). We examined SIRT6 protein expression and mRNA levels in the context of PKCδ depletion and PKCδ overexpression. SIRT6 protein was increased in both shδ110 and shδ680 cells compared to the shNT (Fig. 6A), with similar changes seen in SIRT6 mRNA abundance (Fig. 6B). Conversely, expression of pPKCδFLAG resulted in a dramatic decrease of both SIRT6 protein and mRNA (Fig. 6, C and D). We then asked if the increase in SIRT6 in PKCδ-depleted cells was associated with displacement of KDM2A from chromatin. In the absence of PKCδ, less KDM2A was associated with chromatin in PKCδ-depleted shδ110 and shδ680 cells compared to shNT cells (Fig. 6E). The demonstration that depletion of PKCδ enriches H3K36me2 and reduces KDM2A on chromatin suggests that PKCδ may prime cells for DSB repair through SIRT6. To determine if displacement of KDM2A from chromatin is associated with increased mono-ADP ribosylation (MAR) on KDM2A, we immunoprecipitated KDM2A from nuclear fractions of ParC5 shNT and PKCδ-depleted shδ110 and shδ680 cells and immunoblotted with an anti-MAR antibody. Indeed, we observed more MAR of KDM2A in the PKCδ-depleted shδ110 and shδ680 cells compared to the shNT (Fig. 6F).

**Fig. 6.**
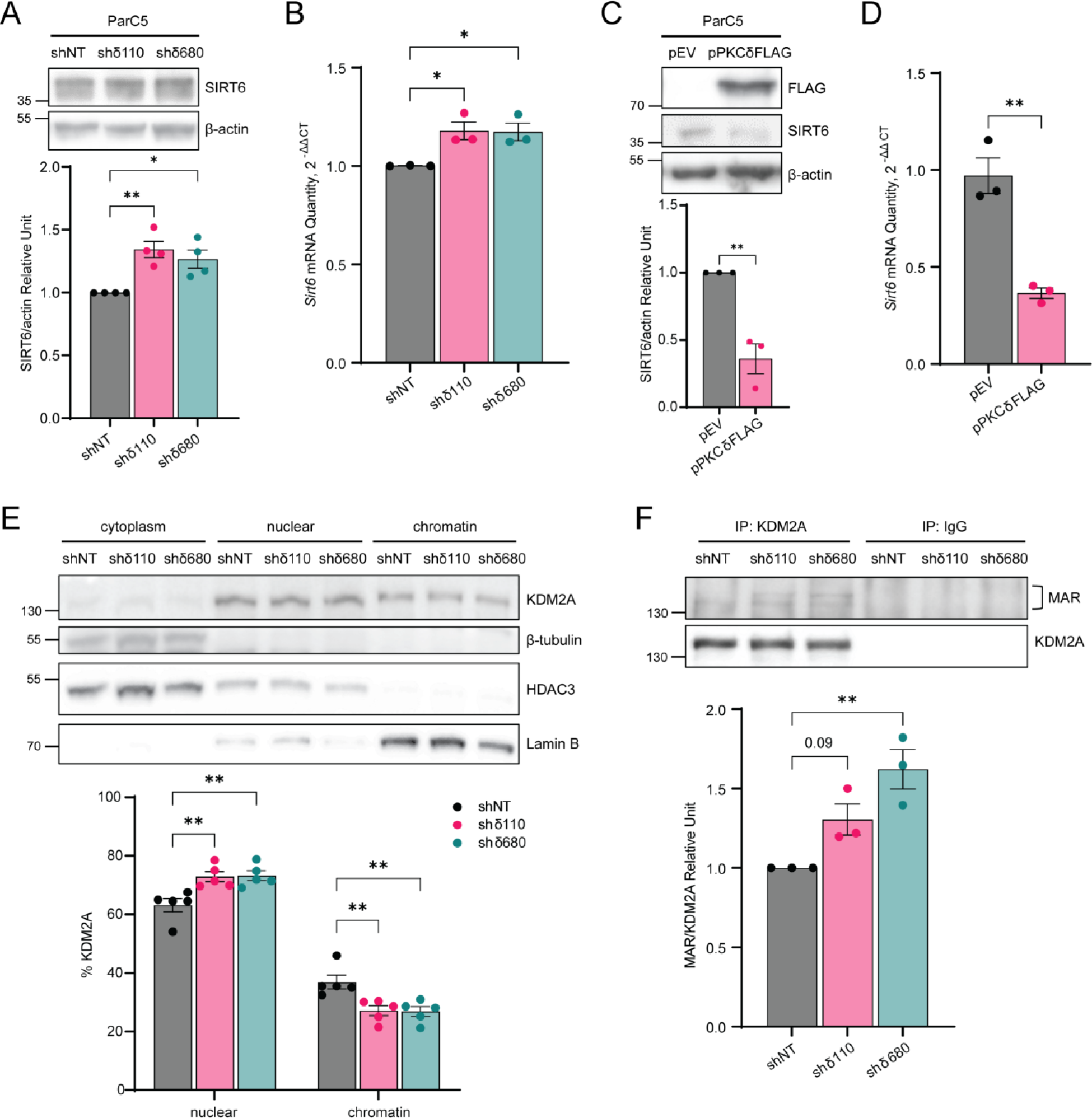
SIRT6 regulates KDM2A ribosylation in PKCδ-depleted cell. (**A**) Top: Whole cell lysates of ParC5 shNT, shδ110, and shδ680 cells were immunoblotted with the indicated antibodies. Bottom: Densitometry of SIRT6 expression normalized to β-actin. (**B**) Relative expression of SIRT6 mRNA in ParC5 shNT, shδ110, and shδ680 cells was assayed by qRT-PCR. (**C**) Top: ParC5 cells transfected with pEV or pPKCδFLAG were analyzed for SIRT6 expression by immunoblot analysis. Bottom: Densitometry of SIRT6 expression normalized to β-actin. (**D**) Relative expression of SIRT6 mRNA in ParC5 overexpressing either pEV or pPKCδFLAG cells was assayed by qRT-PCR. Shown is data from at least three independent biological replicates. Data is shown as mean ± SEM. (**E**) Top: ParC5 shNT, shδ110, and shδ680 cells were fractionated into cytoplasm, nuclear and chromatin fractions, and each fraction was immunoblotted with the indicated antibodies. Bottom: Percentage of KDM2A in nuclear and chromatin were calculated. (**F**) Top: Immunoprecipitation of KDM2A from ParC5 shNT, shδ110, and shδ680 nuclear fractions using KDM2A antibody or normal IgG control, and immunoblotted using anti-mono-ADP-ribose (MAR) and KDM2A antibodies. Bottom: Densitometry of MAR-KDM2A normalized to KDM2A. Statistics represent one-way ANOVA followed by Dunnett’s multiple comparisons for (A), (B) and (F), unpaired t-test for (C) and (D), and two-way ANOVA followed by Dunnett’s multiple comparisons for (E). \**P* < 0.05, \*\**P* < 0.01, \*\*\**P* < 0.001, and \*\*\*\**P* < 0.0001.

To further investigate the role of SIRT6 in PKCδ-dependent regulation of DNA repair and apoptosis, we depleted SIRT6 in ParC5 shNT and shδ110 NHEJ reporter cells using four unique shRNAs (Fig. 7A). shSIRT6-2 and shSIRT6-3 showed approximately 90% reduction of SIRT6 protein levels, and thus these cell lines were used in the remainder of the experiments. We hypothesized that SIRT6 is required to facilitate displacement of chromatin KDM2A and enrich H3K36me2 in PKCδ-depleted cells. When SIRT6 was depleted in shδ110 cells, the abundance of H3K36me2 on chromatin was reduced compared to the shδ110-shSCR control, while no change in H3K36me2 was observed in the shNT cells (Fig. 7B). To determine if the KDM2A/H3K36me2 switch is associated with changes in chromatin structure, we performed an MNase assay. With depletion of SIRT6, the more relaxed chromatin structure seen in the shδ110 cells was reversed, while SIRT6 depletion had no effect on MNase sensitivity in shNT control cells (Fig. 7C). We next examined the ability of SIRT6-depleted cells to repair DNA DSBs. As expected, NHEJ-mediated repair was increased dramatically in shδ110 cells (Fig.7D). Remarkably, both shRNAs targeting SIRT6 completely reversed the increase in NHEJ repair in PKCδ-depleted cells (Fig. 7D). Likewise, depletion of SIRT6 reversed the increased expression of NHEJ repair proteins (Fig. 7, E to H) we observed in PKCδ-depleted cells (Fig. 3, D to G). DNA Lig4, XLF, and XRCC4 expressions were significantly reduced in the shδ110-shSIRT6-2/3 cells compared to shδ110 cells (Fig. 7, E to H).

**Fig. 7.**
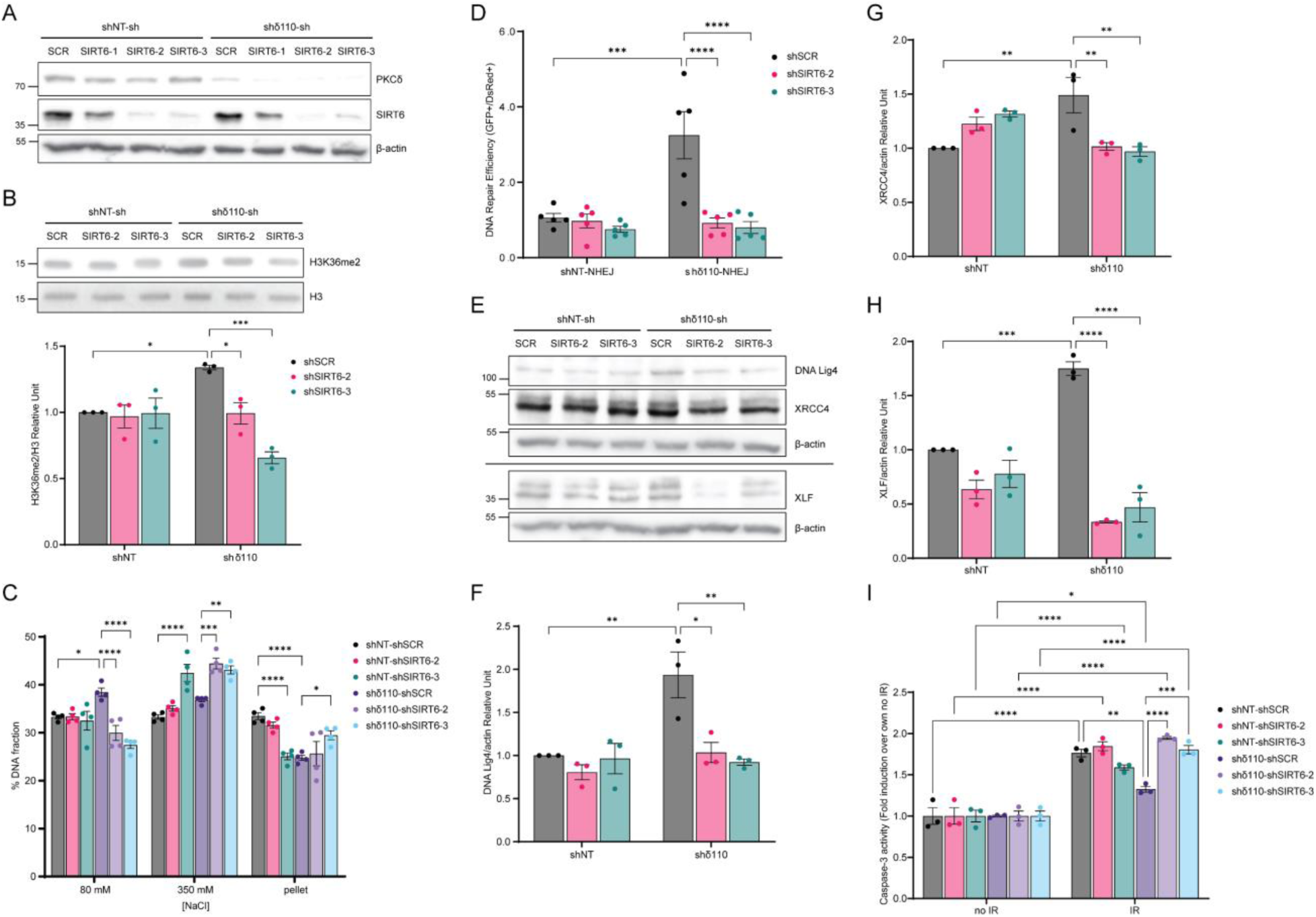
SIRT6 functions downstream of PKCδ to regulate DNA repair and apoptosis. (**A**) ParC5 shNT and shδ110 NHEJ reporter cells were generated that stably express shSCR, shSIRT6- 1, shSIRT6-2, or shSIRT6-3. Lysates were immunoblotted for the indicated proteins. (**B**) Top: Histone fractions from ParC5 shNT/shδ110-shSCR/shSIRT6-2/shSIRT6-3 cells were acid-extracted and immunoblotted for H3K36me2 and total H3. Bottom: Densitometry of H3K36me2 normalized to total H3. (**C**) MNase assay was performed on ParC5 shNT/shδ110-shSCR/shSIRT6- 2/shSIRT6-3 cells. Percentage of DNA in each [NaCl] fraction was assayed as described in Materials and Methods. (**D**) DNA repair was analyzed in ParC5 shNT/shδ110-shSCR/shSIRT6- 2/shSIRT6-3 NHEJ reporter cells and quantified as described in Materials and Methods. (**E to H**) Lysates from ParC5 shNT/shδ110-shSCR/shSIRT6-2/shSIRT6-3 cells were immunoblotted with the indicated antibodies. Densitometry of DNA Lig4 (F), XRCC4 (G), and XLF (H) normalized to their corresponding β-actin control from the Western blots shown in (E). (**I**) ParC5 shNT/shδ110-shSCR/shSIRT6-2/shSIRT6-3 cells were IR at 10 Gy and allowed to recover for 48 hrs. Cells were harvested and caspase-3 activity was assayed. Shown is the fold induction of caspase-3 activities in each cell line over their own corresponding no IR controls. Shown is data from at least three independent biological replicates. Data is shown as mean ± SEM. Statistics represent two-way ANOVA followed by Tukey’s multiple comparisons. \**P* < 0.05, \*\**P* < 0.01, \*\*\**P* < 0.001, and \*\*\*\**P* < 0.0001.

Finally, to determine if SIRT6 is required for radioprotection in PKCδ-depleted cells, caspase-3 activity was assayed in SIRT6-depleted cells following IR. As expected, PKCδ-depleted shδ110 cells showed protection against IR compared to shNT cells (Fig. 7I, gray bar vs dark purple bar), however, this protection was completely reversed when SIRT6 was depleted (shδ110- shSIRT6-2 and shδ110-shSIRT6-3) indicating that SIRT6 is required for radioprotection (Fig. 7I). Taken together, our data identifies SIRT6 a key regulator of cell death downstream of PKCδ (Fig. 8). We propose that in the presence of PKCδ, SIRT6 decreases in abundance, allowing KDM2A to demethylate H3K36me2 on chromatin, which in turn causes chromatin compaction and suppression of NHEJ and HR repair, leading to genomic instability and apoptosis (Fig. 8, left panel). Contrarily, when PKCδ is depleted, cell survival is enhanced through increased abundance of SIRT6, KDM2A disassociation by SIRT6-dependent ribosylation, increased H3K36me2, chromatin relaxation and increased DSB repair efficiency (Fig. 8, right panel).

**Fig. 8.**
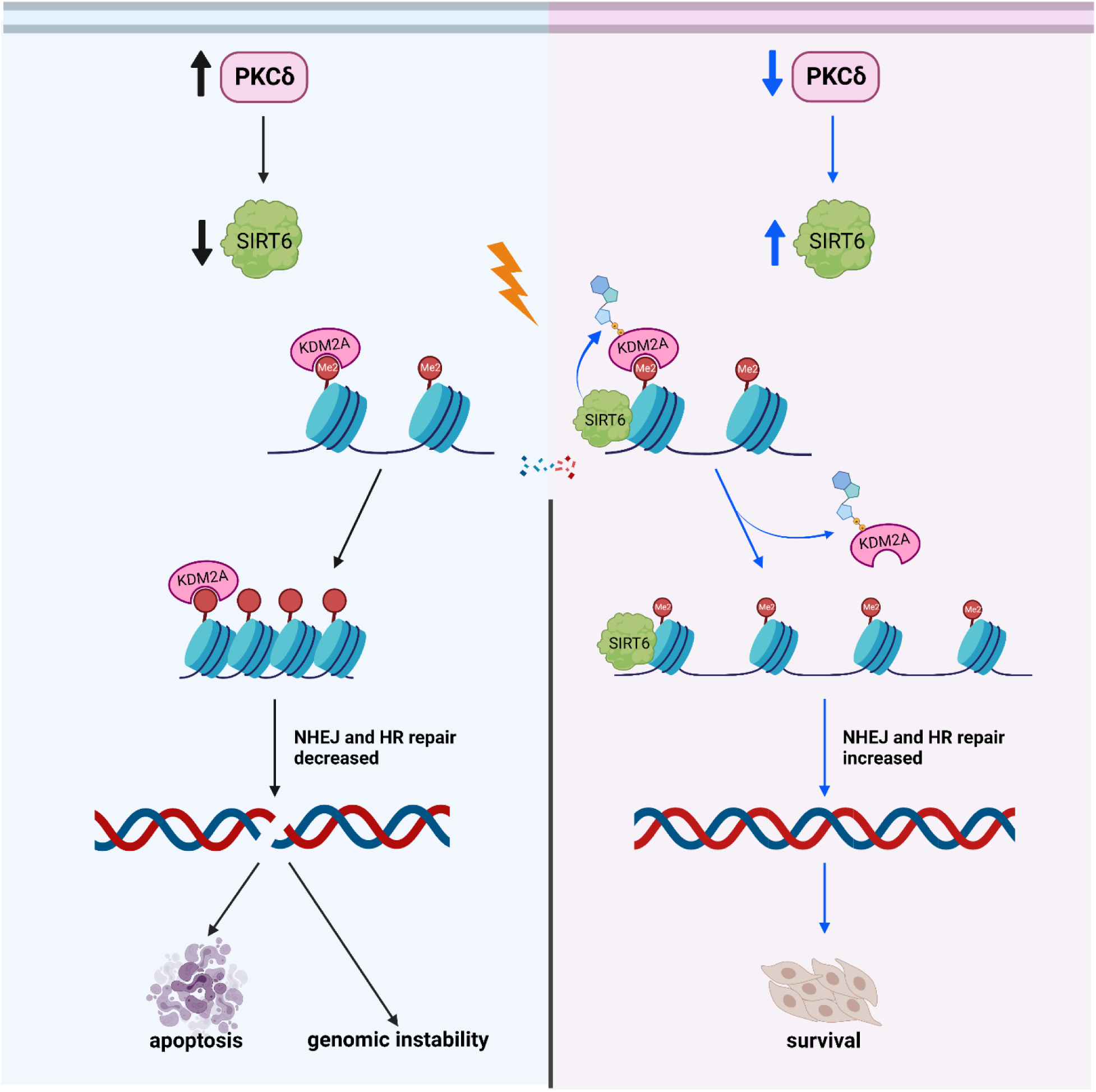
Model of PKCδ-dependent apoptosis. Left, when PKCδ is overexpressed, SIRT6 expression decreases, allowing KDM2A to demethylate H3K36, resulting in more condensed chromatin and inhibition of NHEJ and HR repair. This results in increased genomic instability and apoptosis. Right, depletion of PKCδ increases the expression of SIRT6, which in turn leads to the displacement of KDM2A from chromatin. Accumulation of H3K36me2 results in a more open chromatin structure, increases NHEJ and HR repair, and promotes cell survival.

## DISCUSSION

PKCδ regulates many cellular functions including growth factor signaling at the plasma membrane and apoptosis in the nucleus (*44*). Inhibition of PKCδ, or its nuclear exclusion, provides radioprotection *in vitro* and *in vivo* (*19–26, 29*). Here we show that PKCδ regulates DSB repair and apoptosis through a mechanism that depends on reciprocal regulation of SIRT6. When expression of PKCδ is inhibited, increased expression of SIRT6 regulates histone methylation and KDM2A ribosylation, generating a more open chromatin structure and increasing the accessibility of repair factors to sites of DNA damage. Our data suggests a novel mechanism for control of apoptosis by PKCδ that is dependent on SIRT6 and mediated through regulation of chromatin accessibility and DNA repair.

Previous studies from our lab show that translocation of PKCδ to the nucleus in response to DNA-damaging agents is both necessary and sufficient to induce apoptosis (*22*). Moreover, we show that chromatin changes mediated by PKCδ require both its nuclear localization and kinase function. Nuclear translocation requires progressive tyrosine phosphorylation at Tyr^155^ and Tyr^64^ in the regulatory domain of the kinase, with the initial phosphorylation event at Tyr^155^ mediated by the DNA damage-induced tyrosine kinase, c-Abl (*23, 24*). This links nuclear translocation of PKCδ to the DNA damage response (*45*), and supports previous studies from our lab that show the steady state level of nuclear PKCδ determines responsiveness to DNA damaging agents in a panel of non-small cell lung cancer (NSCLC) cells (*46*). Thus, the cytoplasmic:nuclear partitioning of PKCδ may underly radiosensitivity of both tumors and adjacent normal tissues.

We propose that depletion of PKCδ “primes” cells for DNA repair. Depletion of PKCδ increases NHEJ and HR mediated repair, increases the basal expression of DNA repair proteins and the number of NHEJ and HR foci, and decreases the steady state level of DNA damage. We have previously shown that the tyrosine kinase inhibitors (TKIs), imatinib and dasatinib, which inhibit tyrosine phosphorylation and nuclear translocation of PKCδ, also increase expression of DNA repair proteins and enhance NHEJ and HR repair of DSBs (*25, 47*). Similar regulation of NHEJ and HR RNA transcripts and protein expression can occur in response to hypoxia (decreased) (*48, 49*), H3K9 demethylation (increased) (*50*), reprogramming of iPS cells (increased) (*51*) and in some cancer cells (increased or decreased) (*52, 53*). In most cases, both NHEJ and HR related genes are regulated as a set, reflecting overall activation or suppression of DSB repair. Priming in the case of PKCδ depletion is transcriptional or post-transcriptional, as overexpression or depletion of PKCδ decreases or increases mRNA and protein expression of NHEJ and HR repair factors, respectively. With both depletion of PKCδ and TKIs pretreatment, NHEJ and HR foci form more rapidly after IR (*25, 29, 47*). Thus, depletion of PKCδ may reset the homeostatic environment to respond more rapidly and robustly to DNA damage. Other studies support a role for PKCδ regulation of cell cycle arrest, specifically in mediating G1 arrest through induction of p21 (*54–56*), in S phase arrest (*57*), and in the maintenance of G2/M DNA damage checkpoint (*58*). It seems likely that regulation of checkpoint activation is linked to PKCδ via DNA repair, however we have not observed consistent differences in the cell cycle arrest of PKCδ depleted ParC5 cells basally or after IR (Reyland *et al.*,unpublished data).

Our data indicates a role for PKCδ upstream of DSBs repair pathways. Like transcription and replication, the DNA repair machinery require access to genomic DNA packaged in chromatin. Thus, the ability to dynamically change chromatin structure is essential for the DDR, DNA repair and maintenance of genome integrity. Typically, more open chromatin leads to enhanced DDR and DNA repair, while more compact chromatin inhibits DNA repair (*32*). Our studies demonstrate that in the absence of PKCδ, chromatin is more sensitive to MNase, indicating a more open conformation. Chromatin remodeling can be achieved by histone post-translational modifications such as phosphorylation, and chromatin remodeling enzymes (*32, 59*). While it is widely known that acetylation and ubiquitylation promote an open chromatin structure, the role of methylation in DDR and DNA repair is debated (*60, 61*). While our epiproteomic histone modification analysis identified changes in both acetylation and methylation with PKCδ depletion, most of the changes have not been previously associated with DNA repair, or the change is in the wrong direction. For example, acetylation of H3 on residues Lys^23^ and Lys^14^, and dimethylation on Lys^79^ is decreased. H3K23ac coexists with H3K14ac and is associated with active gene transcription (*62*), however a role in DDR or DNA repair has not been described. H3K79me2 plays a key role in DDR in yeast (*60*), and is linked to active transcription, genomic stability and HR repair (*41*). Since H3K79me2 is reduced in the PKCδ-depleted cells, this histone modification is unlikely to be involved in the phenotypes we show in PKCδ-depleted cells. Conversely, H3K36me2 is typically associated with active open euchromatic regions, is induced upon IR, and accumulates around DSBs (*40*).

Accumulation of H3K36me2 stimulates DSB repair through increased recruitment of the DNA damage sensing proteins Ku70 and Nbs1 (*40*). H3K36me2 is mainly demethylated by KDM2A, a Jumonji demethylase that has been shown to regulate DNA repair through its chromatin-binding capacity, and when the binding capacity of KDM2A to chromatin is disrupted, DNA repair and cell survival are enhanced (*63–65*). Recently, KDM2A has been identified as a substrate of SIRT6, and SIRT6-dependent ribosylation of KDM2A was shown to displace KDM2A from chromatin, increasing H3K36me2 and enhancing NHEJ (*18*). SIRT6 is a NAD^+^- dependent chromatin regulating enzyme that belongs to the Sirtuin family. While it is classified as Class III HDAC catalyzing the deacetylation of Lys^9^, Lys^18^, and Lys^56^ of H3, it has a wide range of substrates other than histones, such as Lys and Arg residues of chromatin regulating enzymes and DNA repair proteins, in which SIRT6 catalyzes the mono-ADP ribosylation of these residues (*66*). We show that PKCδ negatively regulates SIRT6, and that SIRT6 is required for suppression of apoptosis in PKCδ-depleted cells. Mechanistically, SIRT6 is required for all aspects of the radioprotection phenotype we describe in PKCδ-depleted cells, including chromatin relaxation, ribosylation of KDM2A and its dissociation from chromatin, increased H3K36me2 and priming of DNA repair.

While SIRT6 has been identified as one of the earliest DNA damage sensors recruited to DSBs to open chromatin and initiate repair (*13*), little is known about how SIRT6 is regulated in context of DDR and DNA repair. Proteomic analysis has identified conserved residues on SIRT6 that are phosphorylated (Ser^10^, Thr^294^, Ser^303^, Ser^330^, and Ser^338^) (*67*). There are no PKCδ-specific consensus phosphorylation sites in SIRT6, suggesting that PKCδ regulation of SIRT6 is indirect or not mediated by PKCδ kinase activity. In response to oxidative stress, SIRT6 is phosphorylated by c-jun N-terminal kinase (JNK) on Ser^10^ to stimulate DNA repair by recruiting PARP1 to DSBs and through mono-ADP ribosylation of PARP1 on Lys^521^ (*68, 69*). While PKCδ is an upstream effector JNK (*19*), it represses SIRT6, thus it is unlikely that the increased abundance of SIRT6 in PKCδ-depleted cells is through the activation of JNK. SIRT6 protein stability is regulated through phosphorylation of SIRT6 on Ser^338^ by Akt which allows its ubiquitination by MDM2, promoting SIRT6 degradation (*70*). While we have not examined degradation of SIRT6 protein, in our studies PKCδ reciprocally regulates the abundance of SIRT6 mRNA, suggesting that PKCδ may be a negative regulator of SIRT6 transcription.

We show that the catalytic activity of PKCδ is required and sufficient to induce apoptosis and increase DNA damage, however substrate(s) of PKCδ in response to DNA damage are still unknown. Phosphorylation of histone H3 on Ser^10^ is dynamically regulated during the cell cycle, and is essential for mitosis where it regulates chromatin condensation (*71*). Park *et al.* have reported that PKCδ phosphorylation of H3 on Ser^10^ in vitro and in HeLa and Jurkat cells is associated with apoptosis and nuclear condensation, however the significance of these findings to DNA repair is not clear as the authors did not examine chromatin associated histones (*72*). We have previously identified MSK1, a chromatin-modifying kinase, as a downstream effector of PKCδ that is required for apoptosis (*30*). MSK1/2 phosphorylates H3 histones at multiple sites, including Ser^10^ and Ser^28^ (*73*). This raises the possibility that chromatin modification by MSK1 could be a component of the DNA repair program regulated by PKCδ.

Our studies provide a mechanistic understanding of how PKCδ regulates DNA damage and cell death, and have implications for other processes associated with altered DNA repair, including cancer and aging. For example, a reduction in H3K36me2, as well as H3K36M mutations which prevent H3K36 methylation, are associated with brain and bone cancers (*74*). Therefore, the ability of PKCδ to decrease H3K36me2 may explain in part its role as a tumor promoter in mouse models of cancer (*26*). While little is known regarding PKCδ and aging, PKCδ has been implicated in neurodegenerative diseases, including Alzheimer’s and Parkinson’s disease (*26*). Many studies link SIRT6, DNA repair and aging. For example, the Gorbunova group has shown that long-lived species have higher SIRT6 activity and more robust DNA repair activity, suggesting that these activities coevolved (*75*). Further, mice overexpressing *Sirt6* have a longer lifespan (*76*), while mice lacking *Sirt6* display increased genomic instability and a degenerative aging-like phenotype (*77*). We similarly show that overexpression of PKCδ decreases SIRT6 expression and increases genomic instability. Taken together, our findings support further investigation of SIRT6 as a therapeutic target for radioprotection and of PKCδ as a possible target for aging.

## MATERIALS AND METHODS

### Cell culture and generation of expression plasmids

The ParC5 cell line is a rat salivary acinar cell line that has been previously described (*78*). HEK293 cells were obtained from the University of Colorado Cancer Center (UCCC) Cell Technologies Shared Resource, and the human retinal pigmented epithelial (RPE) cell line has been previously described (*79*). The human cell lines were cultured in DMEM/High glucose medium (Cat # SH30243.02, Thermo Fisher Scientific, Logan, UT) supplemented with 10% FBS (Cat # F2442, Sigma, St Louis, MO). Cell line profiling for authentication was done through the UCCC Cell Technologies Shared Resource. Cells used in these experiments were within 10 passages of authentication and were monitored for mycoplasma once a month.

Unless stated otherwise, all PKCδ constructs were constructed in pEGFP-N1 with GFP fused to the C-terminus of PKCδ and have been previously described (*21*). Plasmids used here include pEGFP-PKCδ wild-type (pPKCδ), pEGFP-PKCδ catalytic fragment (pPKCδCF), pEGFP- PKCδCF kinase negative with K376R mutation (pPKCδCFKN), pEGFP-PKCδCF targeted to the cytoplasm (pPKCδCFNLM), pEGFP-PKCδ fused with SV40-NLS sequence to target to the nucleus (pPKCδNLS), and pEGFP-PKCδ targeted to the cytoplasm (pPKCδNLM). To generate pPKCδFLAG, which contains a 2xFLAG tag at the C-terminus, the wild-type mouse *prkcd* gene was amplified using pEGFP-PKCδ as a template. The amplified product was digested with HindIII and BamHI and ligated into pcDNA3.1/2xCTFLAG (a gift of CJ Hu, University of Colorado AMC). The correct final product was verified by DNA sequencing.

For IR experiments, subconfluent cells were irradiated using a Cesium-137 source. For transient expression experiments subconfluent ParC5 cells were transfected using a 2µg:3µL DNA:jetPRIME transfection reagent ratio (Polyplus, New York, NY). For HEK293 and RPE cells, subconfluent cells were transfected using a 1µg:2µL and 1µg:1.5µL DNA:jetPRIME ratio respectively. For inhibitor experiments, cells were treated with 100 nM trichostatin A (TSA, CAS # 58880-19-6, Cayman Chemical, Ann Arbor, MI), 5 mM nicotinamide (NAM, CAS # 98-92-0, Cayman Chemical, Ann Arbor, MI), or 5 mM dimethyl succinate (SUC, CAS # 150731000, Thermo Fisher Scientific, New Jersey, USA) for the indicated times.

### Generation of stable shRNA cell lines

Lentiviruses encoding the appropriate shRNAs were generated as previously described (*80*). ParC5 cells were transduced with lentiviruses encoding unique “non-targeting” shRNAs (Cat # VSC11722, Dharmacon, Lafayette, CO) to generate the shNT control cell line, or two unique species-specific rat shRNAs that target PKCδ (Cat # V3SR11242-243761456, or Cat # V3SR11242-242319029, Dharmacon, Lafayette, CO) to generate the shδ110 and shδ680 cell lines, respectively. HEK293 cells were transduced with lentiviruses that express a non-targeting “scrambled” shRNA (shSCR) as previously described (*29*), or three unique shRNAs targeting human PKCδ (Cat # TRCN0000010193, Cat # TRCN0000010203 or Cat # TRCN0000284800, Open Biosystems, Pittsburg, PA), to generate the shδ193, shδ203 and shδ800 cell lines, respectively. Lentivirus transduced cells were selected in 2 μg/ml puromycin and maintained in media with 1 μg/mL puromycin. To make the rat shSIRT6 cell lines, ParC5 shNT and shδ110 cells containing a stably integrated NHEJ reporter were transduced with lentivirus that express shSCR control (Cat # SHHCTR001, Creative Biogene, Shirley, NY) or three unique shRNAs targeting rat SIRT6 (Cat # SHH407796-1, Cat # SHH407796-2 or Cat # SHH407796-3, Shirley, NY), to generate the stable cell lines referred as shNT-/shδ110-shSCR/shSIRT6-1/shSIRT6-2/shSIRT6-3. For stable expression cells were selected in ParC5 media containing 1 μg/ml puromycin and 0.5 mg/mL hygromycin B, and cell lines were maintained in 1 μg/ml puromycin and 0.4 mg/mL hygromycin B ParC5 media.

### Neutral Comet assay

Subconfluent cells were exposed to 5 Gy IR and harvested at the indicated times for measurement of DNA damage using the neutral comet assay according to the Trevigen Comet Assay protocol (Gaithersburg, MD). Images were analyzed by Trevigen Comet Analysis Software (Version 1.3d). Mean tail moment was used to quantify DNA damage. At least 150 comets were counted for each treatment and each cell line.

### Micronuclei assay

RPE cells were transfected with pEGFP-N1 or pEGFP-PKCδ using jetPRIME as described above for 24 hr, selected and maintained in 1.2 mg/mL geneticin (Life Technologies, Grand Island, NY). Cells were plated on coverslips at passage 3. When cells reached 70-80% confluency they were fixed in 2% paraformaldehyde in PBS for 10 min at room temperature, followed by cell permeabilization with 0.5% Triton X-100 in PBS for 10 min. Cells were washed with PBS twice prior to mounting on slides using Vectashield Vibrance mounting media containing DAPI stain (Cat # H-1800, Vector Laboratories, Burlingame, CA). Images were acquired on an Olympus BX51 fluorescent microscope with a 40x objective. JQuantPlus software (kindly provided by Pavel Lobachevsky, Peter MacCallum Institute, Melbourne, AU) (*81*) was used to identify objects and count micronuclei in each field using the DAPI images. Data is expressed as the average % micronuclei from 20 fields (number of micronuclei / GFP+ cells).

### NHEJ and HR repair assays

The NHEJ and HR reporter plasmids, pCBASceI (pI-SceI expressing plasmid), and pDsRed2-N1 (pDsRed) were kindly provided by Dr. Vera Gorbunova (University of Rochester, NY) (*34, 35*). To generate reporter cell lines, the linearized NHEJ or HR reporter cassettes (0.5 µg) were transfected into parental ParC5 cells, ParC5 shδ cell lines (shNT, shδ110, and sh680), or HEK293 shδ cell lines (shSCR, shδ193, shδ203, and sh800) cells using Amaxa Nucleofector 2B (Basic Nucleofector Kit Primary Mammalian Epithelial Cells, Cat # VPI-1005, Program T-020 for ParC5 cells, and Cell Line Nucleofector Kit V, Cat # VCA-1003, Program Q-001 for HEK293 cells, Lonza, Walkersville, MD). Post 24 hr transfection, cells were selected with 100 µg/mL G418 for 7-10 days and maintained in 100 µg/mL G418 ParC5 media.

DNA repair assays were performed as previously described (*82*). ParC5 or HEK293 reporter cells were co-transfected with 5 µg pI-SceI and 0.1 µg pDsRed using Amaxa Nucleofector 2b. Cells were allowed to repair for 72-96 hrs and were analyzed for GFP and DsRed fluorescence by FACS (Gallios 561). The repair efficiency was calculated as the ratio of GFP+ cells to pDsRed+ cells. For pPKCδFLAG reconstitution, cells were transfected with three plasmids, 5 µg pI-SceI, 0.1 µg pDsRed, and 0.1-1 µg pPKCδFLAG or pEV. For DNA repair assays with inhibitors, cells were pretreated with 100 nM TSA, 5 mM NAM, or 5 mM SUC for 24 hr prior to electroporation and cells were recovered in media with the corresponding inhibitors for 72 hr.

### Analysis and quantification of DNA damage foci

ParC5 cells were grown on coverslips and exposed to 1 Gy IR. Cells were permeabilized prior to fixing in 3% paraformaldehyde in PBS at the indicated time post IR. Foci were detected by immunostaining at a dilution of 1:1000 with anti-γH2AX (Cat # 05-636, RRID:AB_309864, Millipore, Burlington, MA), 1:200 anti-DNA-PK pSer^2056^ (Cat # ab18192, RRID:AB_869495, Abcam, Cambridge, United Kingdom), or anti-Rad51 (Cat # ABE257, RRID:AB_10850319, Millipore), followed by staining with secondary antibody Alexafluor 488 anti-Rabbit or Alexafluor 568 anti-Mouse at dilution of 1:1000. Coverslips were mounted on slides using Vectashield Vibrance mounting media. A minimum of 5 fields per condition were obtained on an Olympus BX51 fluorescent microscope with a 40x objective. JQuantPlus software (*81*) was used to identify objects using the DAPI images, and to count foci. Data is expressed as the average number of foci per cell.

### Immunoblot Analysis

Immunoblot analysis was done as previously described (Ohm 2019). Whole cell lysates were obtained by lysing cell pellets in JNK Lysis buffer (25 mM HEPES pH 7.5, 300 mM NaCl, 1.5 mM MgCl2, 0.2 mM EDTA, 0.1% Triton X-100, 0.5 mM DTT, and 1X HALT Protease and Phosphatase Inhibitor Cocktail (Thermo Fisher Scientific)). Antibodies for PKCδ (Cat # 14188-1- AP, RRID:AB_10638614) and NBS1 (Cat # 55025-1-AP, RRID:AB_10858928) were obtained from Proteintech (Rosemont, IL). Antibodies for DNA Ligase IV (Cat # sc-271299, RRID:AB_10610371), XRCC4 (Cat # sc-271087, RRID:AB_10612396), XLF (Cat # sc-166488, RRID:AB_2152940), SIRT6 (Cat # sc-517556, RRID:AB_2915920), and Mre11 (Cat # sc-135992, RRID:AB_2145244) were purchased from Santa Cruz Biotechnology (Dallas, TX). Antibodies for Artemis (Cat # NBP2-44276, RRID:AB_2915919), β-tubulin (Cat # NB600-936H, RRID:AB_1878245), and H3 (Cat # NBP2-36468, RRID:AB_2925191) were purchased from Novus Biologicals (Centennial, CO). Rad50 (Cat # MA5-35412, RRID:AB_2849313), Rad51 (Cat # ABE257. RRID:AB_10850319), and H3K36me2 (Cat # MA5-14867, RRID:AB_10983670) were purchased from Thermo Fisher Scientific and Millipore respectively. Antibodies for FLAG-tag (Cat # F3165, RRID:AB_259529) was obtained from Sigma-Aldrich. KDM2A (Cat # ab191387, RRID:AB_2928955), Lamin B1 (Cat # ab194109, RRID:AB_2933958), and β-actin (Cat # ab49900, RRID:AB_867494) were purchased from Abcam. HDAC3 (Cat # 85057, RRID:AB_2800047) was purchased from Cell Signaling Technology (Beverley MA). Chemiluminescent images were captured digitally using the KwikQuant Image Analyzer (Kindle Biosciences, Greenwich, CT) and densitometry performed using the KwikQuant analysis software. Immunoblots of total protein lysates were normalized to intensity of β-actin control, unless otherwise indicated.

### qRT-PCR

ParC5 shNT, shδ110, or shδ680 cells were plated in 6-well plates and collected at 60-80% confluency. For overexpression experiment, subconfluent ParC5 cells were transfected with 2 µg pEV or pPKCδFLAG using jetPRIME and let grow for the indicated times. Total RNA was purified using the Quick-RNA miniprep kit (Zymo Research) and concentrations were measured with a Nanodrop ND-1000 spectrophotometer (Nanodrop Technologies). cDNA was synthesized using a Verso cDNA synthesis kit (Thermo Fisher Scientific). All qRT-PCR primers were PrimeTime qPCR Primers purchased directly from IDT: Artemis (*Dclre1c*, Rn.PT.58.11997467), DNA Ligase IV (*Lig4*, Rn.PT.58.34706582), XRCC4 (*Xrcc4*, Rn.PT.58.6049573) XLF (*Nhej1*, Rn.PT.58.11756453), Mre11 (*Mre11*, Rn.PT.58.35382852), Rad50 (*Rad50*, Rn.PT.58.9875506), Nbs1 (*Nbs1*, Rn.PT.58.18317133), Rad51 (*Rad51*, Rn.PT.58.11260740), BRCA1 (*Brca1*, Rn.PT.58.8533844), SIRT6 (*Sirt6*, Rn.PT.58.18496502), GAPDH (*Gapdh*, Rn.PT.39a.11180736.g), Tbp (*Tbp*, Rn.PT,39a.22214837), and B2m (*B2m*, Rn.PT.39a.22214841.g). qRT-PCR measurements were done in StepOnePlus Real-Time PCR Systems (Applied Biosystems) using SYBR Select Master Mix. The C_T_ was determined automatically by the instrument. Relative expression fold change was calculated using the 2^-ΔΔCT^ method. The geometric mean (geNorm) normalizations were performed as previously described (*47*). In short, *Gapdh* was determined to be a good reference gene for the ParC5 shNT, shδ110, and shδ680 cells, while geNorm of *Tbp* and *B2m* genes was used for the ParC5 overexpressing pPKCδFLAG cells. The mRNA expression levels of the geNorm genes were used as an internal control to normalize each qRT-PCR reaction (ΔC_T_). The relative expression levels were calculated as fold enrichment over the Ctrl cells (ΔΔC_T_). All samples were analyzed in biological triplicates.

### Micrococcal nuclease assay

For PKCδ reconstitution, ParC5 shNT and shδ110 cells were transfected with pGFP, pPKCδCF, pPKCδCFKN, pPKCδNLS, or pPKCδNLM using jetPRIME for 24hr. For inhibitor experiments, ParC5 shNT and shδ110 cells were treated with 5 mM NAM or 5 mM SUC for 48 hrs before collection and cell counting. Chromatin preparation and salt fractionation are adapted from previously described methods using 2 x 10^6^ cells per reaction (*38, 39*) with the following modifications. Briefly, cells were harvested at 600 *x g* for 10 min at room temperature and washed once with PBS. Cells were resuspended in 1 mL NE1 buffer (20 mM HEPES pH 7.8, 10 mM KCl, 1 mM MgCl_2_, 1 mM CaCl_2_, 0.1% Triton X-100, 20% glycerol) and incubated on ice for 10 min. Nuclei were collected by centrifugation and resuspended in 200 µL NE1 buffer with 10 U micrococcal nuclease (MNase). Reactions were incubated at 37°C for 8 min and stopped by adding 2 mM EGTA. Supernatants labeled as “S1” were collected by centrifugation. Pellets were resuspended in 200 µL 80 mM Triton buffer (10 mM Tris pH 7.4, 2 mM MgCl_2_, 2 mM EGTA, 0.1% Triton X-100, 70 mM NaCl) and incubated at 4°C with gentle shaking for 2 hr. Supernatants were collected and combined with S1 for the “80 mM” fraction. Pellets were resuspended in 200 µL 350 mM Triton buffer (same as 80 mM Triton but containing 340 mM NaCl) and incubated at 4°C overnight. Supernatants were collected for the “350 mM” fraction, and remaining pellets were resuspended in 200 µL 350 mM Triton buffer for the “pellet” fraction. DNA was extracted from each fraction using phenol-chloroform-isoamyl alcohol. DNA concentration was measured using a Nanodrop ND-1000 spectrophotometer, and the % DNA of each fraction was calculated. Data is expressed as the relative % by comparing to the control.

### Epiproteomic Histone Modification Panel (EHMP)

Profiling of histone modification in PKCδ shRNA and control cells was performed by the Northwestern University Proteomics Core. Briefly, frozen pellets of 3 x 10^6^ ParC5 shNT, shδ110, and shδ680 cells were acid-extracted and samples were derivatized via propionylation reaction and digested with trypsin (*83*). Samples were analyzed by liquid chromatography coupled to TSQ Quantum Ultra mass spectrometry.

### Acid extraction of histones

Acid extraction was performed as described (*84*) with modifications. In brief, 5 million cells were collected and washed with PBS. Pellets were lysed in 1 mL hypotonic lysis buffer (10 mM Tris pH 8.0, 1 mM KCl, 1.5 mM MgCl_2_ with freshly added 1 mM DTT and 1X HALT protease) and rotated for 30 min at 4°C. Nuclei were collected at 10,000 x g for 10 min and resuspended in 400 µL 0.4 N H_2_SO_4_, vortexed and rotated at 4°C overnight. The samples were spun at 16,000 x g for 10 min and the supernatant containing histone fraction was transferred in a new tube. Trichloroacetic acid (TCA) was added to the histone fraction drop by drop to a final volume of 25%, with mixing in between drops, and the solution was incubated at 4°C overnight. Histones were collected at 16,000 x g for 10 min and washed with ice cold acetone three times. Finally, histone pellets were air-dried and dissolved in water.

### Cell fractionation

Cell fractionation was performed as described (*85, 86*) with modifications. ParC5 cells were resuspended in hypotonic buffer (10 mM Tris pH 7.9, 10 mM KCl, 0.1 mM EDTA, 0.5 mM EGTA, 1 mM DTT, 1 mM PMSF and 1X HALT protease and incubated on ice for 15 min. Then, added 10% NP-40, vortexed at high speed for 10 sec, centrifuged at top speed for 30 sec, and the supernatant was collected as “cytoplasmic” fraction. The nuclei pellet was resuspended in high salt buffer (20 mM HEPES pH 7.9, 400 mM NaCl, 1 mM EDTA, 5 mM EGTA, 1 mM DTT, 1 mM PMSF, and 1X HALT protease and phosphatase inhibitors), rotated at 4°C for 15 min, centrifuged at top speed for 5 min, and the supernatant was collected as “nuclear” fraction. The pellet was resuspended in MNase buffer (20 mM Tris pH 7.5, 100 mM KCl, 2 mM MgCl_2_, 1 mM CaCl_2_, 300 mM sucrose, 0.1% Triton X-100, 1 mM DTT, 1 mM PMSF, and 1X HALT protease and phosphatase inhibitors), and sonicated using a Bioruptor Pico (Diagenode, Denville, NJ) for 10 min (30 sec ON and 30 sec OFF). Then, added MNase (5 U/2 million cells) for 30 min at room temperature followed by addition of 5 mM EDTA to stop the reactions. Samples were then centrifuged at 2000 x g for 5 min at 4°C and the supernatant was collected as “chromatin” fraction.

### Immunoprecipitation

Nuclear fractions (0.5-1 mg) of ParC5 shNT, shδ110, or shδ680 cells from above were brought up to 1 ml with Low Salt IP Buffer (20 mM HEPES pH 7.9, 150 mM NaCl, 0.2 mM EDTA, 10% glycerol, 0.1% NP-40, 1X HALT Protease and Phosphatase Inhibitor Cocktail). Lysates were precleared with 25 µL Protein A/G PLUS-agarose (Cat # sc-2003, Santa Cruz Biotechnology) at 4°C for 1 hr. The precleared lysates were transferred to new tubes and mixed with Rabbit anti-KDM2A antibody (Cat # ab191387) or Rabbit IgG control (Cat # 3900S, Cell Signaling Technology) for 4 hr and up to overnight at 4°C. The antibody-bound proteins were captured with Dynabeads Protein A (Thermo Fisher Scientific) for 2 hr at 4°C. The immunocomplexes were then washed in Low Salt IP buffer before eluting in 2X SDS buffer at 80°C for 10 min. The eluants were separated by SDS-PAGE and probed with anti-mono-ADP ribose antibody (Cat # AbD33204, Bio-Rad) and anti-KDM2A antibody.

### Caspase-3 activity assay

ParC5 were transfected with 2 µg pEV or pPKCδFLAG using jetPRIME for 72 hrs. Cells were then harvested and assayed for caspase-3 activity with the Caspase-3 Cellular Activity Assay Kit PLUS (Biomol, Farmingdale, NY), which uses N-acetyl-DEVD-p-nitroaniline as a substrate, according to the manufacturer’s instructions.

### Statistical Analysis

Data shown in figures are representative experiments repeated a minimum of three times. Graphical data are presented as mean ± standard error of mean (SEM) unless otherwise stated. DNA damage foci, comet assay and qRT-PCR were designed with a minimum of triplicate biological samples. Statistics were determined using GraphPad Prism 8 software. Equal variance and normality were tested to decide which kind of test was run. A two-way ANOVA (α = 0.05) with Dunnett’s multiple comparisons was performed unless otherwise indicated (\**P* < 0.05, \*\**P* < 0.01, \*\*\**P* < 0.001, and \*\*\*\**P* < 0.0001) within each time point or treatment, comparing each column with the corresponding control.

## Supplementary Materials

TCGA Analysis

Fig. S1. Correlation of copy number variants (CNV) and PKCδ expression in LUAD and HNSCC.

Fig. S2. Overexpression and knockdown of PKCδ in ParC5 and HEK293 cells.

Fig. S3. PKCδ regulates expression of genes required for DNA repair.

Data file S1. Results of mass spectrometry histone profiling of shNT, shδ110 and shδ680 cells. Spreadsheet includes average percent of each modification, standard deviation, and coefficient of variance (CV).

## Acknowledgments

We thank Vera Gorbunova (University of Rochester) for providing the pI-SceI and pDsRed plasmids. Proteomics services were performed by the Northwestern Proteomics Core Facility, generously supported by NCI CCSG P30 CA060553 awarded to the Robert H Lurie Comprehensive Cancer Center, instrumentation award (S10OD025194) from NIH Office of Director, and the National Resource for Translational and Developmental Proteomics supported by P41 GM108569. The TCGA results published here are in whole based upon data generated by the TCGA research Network: https://www.cancer.gov/tcga.

## Author contributions

Conceptualization: MER, TA, JCB

Methodology: TA, AMO, AH, JCB, GW

Investigation: TA, AMO, AH, GW

Funding acquisition: MER, TA

Project administration: TA, AMO

Supervision: MER, JCB, AMO

Writing – original draft: MER, TA

Writing – review & editing: MER, TA, JCB

## Data and materials availability

All data are available in the main text or the supplementary materials.

## References and Notes

1. T. Kumagai, F. Rahman, A. M. Smith, The Microbiome and Radiation Induced-Bowel Injury: Evidence for Potential Mechanistic Role in Disease Pathogenesis. Nutrients 10, (2018).

2. S. Costa, M. R. Reagan, Therapeutic Irradiation: Consequences for Bone and Bone Marrow Adipose Tissue. Front Endocrinol (Lausanne) 10, 587 (2019).

3. R. L. Siegel, K. D. Miller, H. E. Fuchs, A. Jemal, Cancer statistics, 2022. CA Cancer J Clin 72, 7–33 (2022).

4. V. Gregoire, J. A. Langendijk, S. Nuyts, Advances in Radiotherapy for Head and Neck Cancer. J Clin Oncol 33, 3277–3284 (2015).

5. S. B. Jensen, A. Vissink, K. H. Limesand, M. E. Reyland, Salivary Gland Hypofunction and Xerostomia in Head and Neck Radiation Patients. J Natl Cancer Inst Monogr 2019, (2019).

6. M. King et al., Use of Amifostine for Cytoprotection during Radiation Therapy: A Review. Oncology 98, 61–80 (2020).

7. T. Nakano et al., Formation of clustered DNA damage in vivo upon irradiation with ionizing radiation: Visualization and analysis with atomic force microscopy. Proc Natl Acad Sci U S A 119, e2119132119 (2022).

8. Z. Mao, M. Bozzella, A. Seluanov, V. Gorbunova, Comparison of nonhomologous end joining and homologous recombination in human cells. DNA Repair (Amst) 7, 1765–1771 (2008).

9. W. L. Santivasi, F. Xia, Ionizing radiation-induced DNA damage, response, and repair. Antioxid Redox Signal 21, 251–259 (2014).

10. R. Ceccaldi, B. Rondinelli, A. D. D’Andrea, Repair Pathway Choices and Consequences at the Double-Strand Break. Trends Cell Biol 26, 52–64 (2016).

11. G. D. Bowman, M. G. Poirier, Post-translational modifications of histones that influence nucleosome dynamics. Chem Rev 115, 2274–2295 (2015).

12. J. Stadler, H. Richly, Regulation of DNA Repair Mechanisms: How the Chromatin Environment Regulates the DNA Damage Response. Int J Mol Sci 18, (2017).

13. L. Onn et al., SIRT6 is a DNA double-strand break sensor. Elife 9, (2020).

14. D. Toiber et al., SIRT6 recruits SNF2H to DNA break sites, preventing genomic instability through chromatin remodeling. Mol Cell 51, 454–468 (2013).

15. D. B. Lombard, B. Schwer, F. W. Alt, R. Mostoslavsky, SIRT6 in DNA repair, metabolism and ageing. J Intern Med 263, 128–141 (2008).

16. G. Liszt, E. Ford, M. Kurtev, L. Guarente, Mouse Sir2 homolog SIRT6 is a nuclear ADP-ribosyltransferase. J Biol Chem 280, 21313–21320 (2005).

17. W. W. Wang, Y. Zeng, B. Wu, A. Deiters, W. R. Liu, A Chemical Biology Approach to Reveal Sirt6-targeted Histone H3 Sites in Nucleosomes. ACS Chem Biol 11, 1973–1981 (2016).

18. S. Rezazadeh et al., SIRT6 mono-ADP ribosylates KDM2A to locally increase H3K36me2 at DNA damage sites to inhibit transcription and promote repair. Aging (Albany NY) 12, 11165–11184 (2020).

19. M. E. Reyland, S. M. Anderson, A. A. Matassa, K. A. Barzen, D. O. Quissell, Protein kinase C delta is essential for etoposide-induced apoptosis in salivary gland acinar cells. J Biol Chem 274, 19115–19123 (1999).

20. M. J. Humphries et al., Suppression of apoptosis in the protein kinase Cdelta null mouse in vivo. J Biol Chem 281, 9728–9737 (2006).

21. T. A. DeVries, M. C. Neville, M. E. Reyland, Nuclear import of PKCdelta is required for apoptosis: identification of a novel nuclear import sequence. EMBO J 21, 6050–6060 (2002).

22. T. A. DeVries-Seimon, A. M. Ohm, M. J. Humphries, M. E. Reyland, Induction of apoptosis is driven by nuclear retention of protein kinase C delta. J Biol Chem 282, 22307–22314 (2007).

23. M. J. Humphries, A. M. Ohm, J. Schaack, T. S. Adwan, M. E. Reyland, Tyrosine phosphorylation regulates nuclear translocation of PKCdelta. Oncogene 27, 3045–3053 (2008).

24. T. S. Adwan, A. M. Ohm, D. N. Jones, M. J. Humphries, M. E. Reyland, Regulated binding of importin-alpha to protein kinase Cdelta in response to apoptotic signals facilitates nuclear import. J Biol Chem 286, 35716–35724 (2011).

25. S. M. Wie, T. S. Adwan, J. DeGregori, S. M. Anderson, M. E. Reyland, Inhibiting tyrosine phosphorylation of protein kinase Cdelta (PKCdelta) protects the salivary gland from radiation damage. J Biol Chem 289, 10900–10908 (2014).

26. J. D. Black, T. Affandi, A. R. Black, M. E. Reyland, PKCalpha and PKCdelta: Friends and Rivals. J Biol Chem 298, 102194 (2022).

27. D. J. Keszenman, L. Kolodiuk, J. E. Baulch, DNA damage in cells exhibiting radiation-induced genomic instability. Mutagenesis 30, 451–458 (2015).

28. M. E. Reyland, K. A. Barzen, S. M. Anderson, D. O. Quissell, A. A. Matassa, Activation of PKC is sufficient to induce an apoptotic program in salivary gland acinar cells. Cell Death Differ 7, 1200–1209 (2000).

29. S. M. Wie, E. Wellberg, S. D. Karam, M. E. Reyland, Tyrosine Kinase Inhibitors Protect the Salivary Gland from Radiation Damage by Inhibiting Activation of Protein Kinase C-delta. Mol Cancer Ther 16, 1989–1998 (2017).

30. A. M. Ohm, T. Affandi, M. E. Reyland, EGF receptor and PKCdelta kinase activate DNA damage-induced pro-survival and pro-apoptotic signaling via biphasic activation of ERK and MSK1 kinases. J Biol Chem 294, 4488–4497 (2019).

31. A. A. Matassa, L. Carpenter, T. J. Biden, M. J. Humphries, M. E. Reyland, PKCdelta is required for mitochondrial-dependent apoptosis in salivary epithelial cells. J Biol Chem 276, 29719–29728 (2001).

32. N. Nair, M. Shoaib, C. S. Sorensen, Chromatin Dynamics in Genome Stability: Roles in Suppressing Endogenous DNA Damage and Facilitating DNA Repair. Int J Mol Sci 18, (2017).

33. M. Fenech et al., Micronuclei as biomarkers of DNA damage, aneuploidy, inducers of chromosomal hypermutation and as sources of pro-inflammatory DNA in humans. Mutat Res Rev Mutat Res 786, 108342 (2020).

34. A. Seluanov, D. Mittelman, O. M. Pereira-Smith, J. H. Wilson, V. Gorbunova, DNA end joining becomes less efficient and more error-prone during cellular senescence. Proc Natl Acad Sci U S A 101, 7624–7629 (2004).

35. Z. Mao, A. Seluanov, Y. Jiang, V. Gorbunova, TRF2 is required for repair of nontelomeric DNA double-strand breaks by homologous recombination. Proc Natl Acad Sci U S A 104, 13068–13073 (2007).

36. L. J. Kuo, L. X. Yang, Gamma-H2AX - a novel biomarker for DNA double-strand breaks. In Vivo 22, 305–309 (2008).

37. M. M. Sanders, Fractionation of nucleosomes by salt elution from micrococcal nuclease-digested nuclei. J Cell Biol 79, 97–109 (1978).

38. S. Henikoff, J. G. Henikoff, A. Sakai, G. B. Loeb, K. Ahmad, Genome-wide profiling of salt fractions maps physical properties of chromatin. Genome Res 19, 460–469 (2009).

39. S. S. Teves, S. Henikoff, Salt fractionation of nucleosomes for genome-wide profiling. Methods Mol Biol 833, 421–432 (2012).

40. S. Fnu et al., Methylation of histone H3 lysine 36 enhances DNA repair by nonhomologous end-joining. Proc Natl Acad Sci U S A 108, 540–545 (2011).

41. M. Ljungman, L. Parks, R. Hulbatte, K. Bedi, The role of H3K79 methylation in transcription and the DNA damage response. Mutat Res Rev Mutat Res 780, 48–54 (2019).

42. V. Kari et al., The histone methyltransferase DOT1L is required for proper DNA damage response, DNA repair, and modulates chemotherapy responsiveness. Clin Epigenetics 11, 4 (2019).

43. M. R. Duan, M. J. Smerdon, Histone H3 lysine 14 (H3K14) acetylation facilitates DNA repair in a positioned nucleosome by stabilizing the binding of the chromatin Remodeler RSC (Remodels Structure of Chromatin). J Biol Chem 289, 8353–8363 (2014).

44. A. X. Wu-Zhang, A. N. Murphy, M. Bachman, A. C. Newton, Isozyme-specific interaction of protein kinase Cdelta with mitochondria dissected using live cell fluorescence imaging. J Biol Chem 287, 37891–37906 (2012).

45. M. E. Reyland, D. N. Jones, Multifunctional roles of PKCdelta: Opportunities for targeted therapy in human disease. Pharmacol Ther 165, 1–13 (2016).

46. A. M. Ohm, A. C. Tan, L. E. Heasley, M. E. Reyland, Co-dependency of PKCdelta and K-Ras: inverse association with cytotoxic drug sensitivity in KRAS mutant lung cancer. Oncogene 36, 4370–4378 (2017).

47. T. Affandi, A. M. Ohm, D. Gaillard, A. Haas, M. E. Reyland, Tyrosine kinase inhibitors protect the salivary gland from radiation damage by increasing DNA double-strand break repair. J Biol Chem 296, 100401 (2021).

48. R. S. Bindra et al., Down-regulation of Rad51 and decreased homologous recombination in hypoxic cancer cells. Mol Cell Biol 24, 8504–8518 (2004).

49. A. R. Kaplan, P. M. Glazer, Impact of hypoxia on DNA repair and genome integrity. Mutagenesis 35, 61–68 (2020).

50. H. Y. Jeon, A. Hussain, J. Qi, Role of H3K9 demethylases in DNA double-strand break repair. J Cancer Biol 1, 10–15 (2020).

51. M. Shimada, K. Tsukada, N. Kagawa, Y. Matsumoto, Reprogramming and differentiation-dependent transcriptional alteration of DNA damage response and apoptosis genes in human induced pluripotent stem cells. J Radiat Res 60, 719–728 (2019).

52. M. Szatkowska, R. Krupa, Regulation of DNA Damage Response and Homologous Recombination Repair by microRNA in Human Cells Exposed to Ionizing Radiation. Cancers (Basel) 12, (2020).

53. C. G. Broustas, H. B. Lieberman, DNA damage response genes and the development of cancer metastasis. Radiat Res 181, 111–130 (2014).

54. K. Yoshida, H. Liu, Y. Miki, Protein kinase C delta regulates Ser46 phosphorylation of p53 tumor suppressor in the apoptotic response to DNA damage. J Biol Chem 281, 5734–5740 (2006).

55. M. Nakagawa et al., Phorbol ester-induced G1 phase arrest selectively mediated by protein kinase Cdelta-dependent induction of p21. J Biol Chem 280, 33926–33934 (2005).

56. G. Perletti et al., p21(Waf1/Cip1) and p53 are downstream effectors of protein kinase C delta in tumor suppression and differentiation in human colon cancer cells. Int J Cancer 113, 42–53 (2005).

57. A. E. Santiago-Walker et al., Protein kinase C delta stimulates apoptosis by initiating G1 phase cell cycle progression and S phase arrest. J Biol Chem 280, 32107–32114 (2005).

58. E. L. LaGory, L. A. Sitailo, M. F. Denning, The protein kinase Cdelta catalytic fragment is critical for maintenance of the G2/M DNA damage checkpoint. J Biol Chem 285, 1879–1887 (2010).

59. B. D. Price, A. D. D’Andrea, Chromatin remodeling at DNA double-strand breaks. Cell 152, 1344–1354 (2013).

60. F. Gong, K. M. Miller, Histone methylation and the DNA damage response. Mutat Res Rev Mutat Res 780, 37–47 (2019).

61. J. L. Miller, P. A. Grant, The role of DNA methylation and histone modifications in transcriptional regulation in humans. Subcell Biochem 61, 289–317 (2013).

62. B. J. Klein et al., Histone H3K23-specific acetylation by MORF is coupled to H3K14 acylation. Nat Commun 10, 4724 (2019).

63. L. L. Cao et al., ATM-mediated KDM2A phosphorylation is required for the DNA damage repair. Oncogene 35, 301–313 (2016).

64. E. Dimitrova, A. H. Turberfield, R. J. Klose, Histone demethylases in chromatin biology and beyond. EMBO Rep 16, 1620–1639 (2015).

65. L. Liu, J. Liu, Q. Lin, Histone demethylase KDM2A: Biological functions and clinical values (Review). Exp Ther Med 22, 723 (2021).

66. F. Fiorentino et al., The Two-Faced Role of SIRT6 in Cancer. Cancers (Basel) 13, (2021).

67. Y. V. Miteva, I. M. Cristea, A proteomic perspective of Sirtuin 6 (SIRT6) phosphorylation and interactions and their dependence on its catalytic activity. Mol Cell Proteomics 13, 168–183 (2014).

68. Z. Mao et al., SIRT6 promotes DNA repair under stress by activating PARP1. Science 332, 1443–1446 (2011).

69. M. Van Meter et al., JNK Phosphorylates SIRT6 to Stimulate DNA Double-Strand Break Repair in Response to Oxidative Stress by Recruiting PARP1 to DNA Breaks. Cell Rep 16, 2641–2650 (2016).

70. U. Thirumurthi et al., MDM2-mediated degradation of SIRT6 phosphorylated by AKT1 promotes tumorigenesis and trastuzumab resistance in breast cancer. Sci Signal 7, ra71 (2014).

71. B. Perez-Cadahia, B. Drobic, J. R. Davie, H3 phosphorylation: dual role in mitosis and interphase. Biochem Cell Biol 87, 695–709 (2009).

72. C. H. Park, K. T. Kim, Apoptotic phosphorylation of histone H3 on Ser-10 by protein kinase Cdelta. PLoS One 7, e44307 (2012).

73. K. M. Reyskens, J. S. Arthur, Emerging Roles of the Mitogen and Stress Activated Kinases MSK1 and MSK2. Front Cell Dev Biol 4, 56 (2016).

74. B. J. Klein et al., Recognition of cancer mutations in histone H3K36 by epigenetic writers and readers. Epigenetics 13, 683–692 (2018).

75. X. Tian et al., SIRT6 Is Responsible for More Efficient DNA Double-Strand Break Repair in Long-Lived Species. Cell 177, 622–638 e622 (2019).

76. Y. Kanfi et al., The sirtuin SIRT6 regulates lifespan in male mice. Nature 483, 218–221 (2012).

77. R. Mostoslavsky et al., Genomic instability and aging-like phenotype in the absence of mammalian SIRT6. Cell 124, 315–329 (2006).

78. D. O. Quissell, K. A. Barzen, R. S. Redman, J. M. Camden, J. T. Turner, Development and characterization of SV40 immortalized rat parotid acinar cell lines. In Vitro Cell Dev Biol Anim 34, 58–67 (1998).

79. J. C. Black et al., KDM4A lysine demethylase induces site-specific copy gain and rereplication of regions amplified in tumors. Cell 154, 541–555 (2013).

80. L. Marek et al., Fibroblast growth factor (FGF) and FGF receptor-mediated autocrine signaling in non-small-cell lung cancer cells. Mol Pharmacol 75, 196–207 (2009).

81. A. N. Ivashkevich et al., gammaH2AX foci as a measure of DNA damage: a computational approach to automatic analysis. Mutat Res 711, 49–60 (2011).

82. A. Seluanov, Z. Mao, V. Gorbunova, Analysis of DNA double-strand break (DSB) repair in mammalian cells. J Vis Exp, (2010).

83. B. A. Garcia et al., Chemical derivatization of histones for facilitated analysis by mass spectrometry. Nat Protoc 2, 933–938 (2007).

84. D. Shechter, H. L. Dormann, C. D. Allis, S. B. Hake, Extraction, purification and analysis of histones. Nat Protoc 2, 1445–1457 (2007).

85. E. Schreiber, P. Matthias, M. M. Muller, W. Schaffner, Rapid detection of octamer binding proteins with ‘mini-extracts’, prepared from a small number of cells. Nucleic Acids Res 17, 6419 (1989).

86. J. Mendez, B. Stillman, Chromatin association of human origin recognition complex, cdc6, and minichromosome maintenance proteins during the cell cycle: assembly of prereplication complexes in late mitosis. Mol Cell Biol 20, 8602–8612 (2000).

