## Supplementary material for "PKCδ regulates chromatin remodeling and DNA repair through SIRT6": Data File S1

| Mark | Sample | Total Average Percent | Total Standard Deviation | Total Percent CV |
|---|---|---|---|---|
| H3.1: K27UN | ParC5_sh110 | 17.3519204881108 | 0.0258871589858689 | 0.149189013421346 |
| H3.1: K27UN | ParC5_sh680 | 16.404965116973 | 0.35940478348493 | 2.1908293063853 |
| H3.1: K27UN | ParC5_shNT | 15.0273598782173 | 0.278471753481303 | 1.8530983202509 |
| H3.1: K27ME1 | ParC5_sh110 | 21.5622242295861 | 0.693688161148782 | 3.2171456606826 |
| H3.1: K27ME1 | ParC5_sh680 | 18.309095986962 | 0.241161021603677 | 1.31716509529148 |
| H3.1: K27ME1 | ParC5_shNT | 16.9032816123654 | 0.996256368163605 | 5.89386363553693 |
| H3.1: K27ME2 | ParC5_sh110 | 47.2426141342158 | 0.719667577039067 | 1.52334410410588 |
| H3.1: K27ME2 | ParC5_sh680 | 50.6245123803261 | 0.410815304171504 | 0.81149483689873 |
| H3.1: K27ME2 | ParC5_shNT | 52.8897220325356 | 0.932891834716797 | 1.7638433307381 |
| H3.1: K27ME3 | ParC5_sh110 | 12.6730005238301 | 0.0537745594423838 | 0.424323816141781 |
| H3.1: K27ME3 | ParC5_sh680 | 13.6269154839434 | 0.15902721317375 | 1.16700814179946 |
| H3.1: K27ME3 | ParC5_shNT | 13.9288413401464 | 0.274367676428106 | 1.96978104443842 |
| H3.1: K27AC | ParC5_sh110 | 1.17024062425721 | 0.0168981758666581 | 1.44399156176824 |
| H3.1: K27AC | ParC5_sh680 | 1.03451103179554 | 0.0200169310494596 | 1.93491711873942 |
| H3.1: K27AC | ParC5_shNT | 1.25079513673526 | 0.0111523135569663 | 0.891617918028947 |
| H3.1: K36UN | ParC5_sh110 | 46.0488259534895 | 0.463840954996789 | 1.00728074037171 |
| H3.1: K36UN | ParC5_sh680 | 51.3705045457942 | 0.690444271790145 | 1.34404806395205 |
| H3.1: K36UN | ParC5_shNT | 52.1540333339706 | 0.964322050081999 | 1.84898844525968 |
| H3.1: K36ME1 | ParC5_sh110 | 12.0539282524205 | 0.0762710346352496 | 0.632748370805461 |
| H3.1: K36ME1 | ParC5_sh680 | 11.4492044225459 | 0.389025117665985 | 3.3978353718614 |
| H3.1: K36ME1 | ParC5_shNT | 12.7443229061274 | 0.141030474088472 | 1.1066140988994 |
| H3.1: K36ME2 | ParC5_sh110 | 26.4098286315609 | 0.161272964385294 | 0.610655096006816 |
| H3.1: K36ME2 | ParC5_sh680 | 23.156950069274 | 0.201857097467168 | 0.871691206585118 |
| H3.1: K36ME2 | ParC5_shNT | 21.9259138013491 | 0.483647305822749 | 2.20582508079089 |
| H3.1: K36ME3 | ParC5_sh110 | 15.335647100149 | 0.253372758081623 | 1.65218172032115 |
| H3.1: K36ME3 | ParC5_sh680 | 13.7609179193733 | 0.115031632337132 | 0.835929935859769 |
| H3.1: K36ME3 | ParC5_shNT | 12.5946394511924 | 0.333512030537406 | 2.6480474636043 |
| H3.1: K36AC | ParC5_sh110 | 0.15177006238014 | 0.0104599917906524 | 6.89199940133985 |
| H3.1: K36AC | ParC5_sh680 | 0.262423043012578 | 0.0312683652126616 | 11.9152513642496 |
| H3.1: K36AC | ParC5_shNT | 0.581090507360463 | 0.0268621088877067 | 4.62270654010934 |

| Mark | Sample | Total Average Percent | Total Standard Deviation | Total Percent CV |
|---|---|---|---|---|
| H3.3: K27UN | ParC5_sh110 | 23.9627464100133 | 0.434923469431334 | 1.81499842292531 |
| H3.3: K27UN | ParC5_sh680 | 23.0579368306526 | 0.363921690105046 | 1.57829251063459 |
| H3.3: K27UN | ParC5_shNT | 21.4863270859555 | 0.252851839453679 | 1.17680345478384 |
| H3.3: K27ME1 | ParC5_sh110 | 27.1973083086591 | 0.621764085154776 | 2.28612360494814 |
| H3.3: K27ME1 | ParC5_sh680 | 24.0973884509286 | 0.382809958357583 | 1.58859520871786 |
| H3.3: K27ME1 | ParC5_shNT | 22.7280791828062 | 1.42274228456326 | 6.25984392750427 |
| H3.3: K27ME2 | ParC5_sh110 | 41.9699911136013 | 0.501946126091558 | 1.19596433731169 |
| H3.3: K27ME2 | ParC5_sh680 | 45.2845396470976 | 0.426356529666862 | 0.941505716938845 |
| H3.3: K27ME2 | ParC5_shNT | 47.3937383935818 | 1.28558127167809 | 2.71255510802284 |
| H3.3: K27ME3 | ParC5_sh110 | 4.2536710379741 | 0.0327034679715843 | 0.768829269579813 |
| H3.3: K27ME3 | ParC5_sh680 | 4.84781234086664 | 0.120984875353466 | 2.49565921381844 |
| H3.3: K27ME3 | ParC5_shNT | 5.18111113750096 | 0.120668237154687 | 2.32900306425177 |
| H3.3: K27AC | ParC5_sh110 | 2.6162831297522 | 0.0292220253399727 | 1.11692901305909 |
| H3.3: K27AC | ParC5_sh680 | 2.71232273045446 | 0.108654726941675 | 4.00596602025564 |
| H3.3: K27AC | ParC5_shNT | 3.21074420015553 | 0.300793382330669 | 9.36833841562645 |
| H3.3: K36UN | ParC5_sh110 | 34.0311482579089 | 0.419030690772881 | 1.23131516925967 |
| H3.3: K36UN | ParC5_sh680 | 38.0695659001023 | 0.564400787072035 | 1.4825511500527 |
| H3.3: K36UN | ParC5_shNT | 39.2963500890201 | 1.39110972490015 | 3.54004817686323 |
| H3.3: K36ME1 | ParC5_sh110 | 7.91139674491804 | 0.220565030652578 | 2.78794045810255 |
| H3.3: K36ME1 | ParC5_sh680 | 7.6987851436022 | 0.277026580125362 | 3.59831551287771 |
| H3.3: K36ME1 | ParC5_shNT | 8.24672608215162 | 0.144072667125414 | 1.7470286473711 |
| H3.3: K36ME2 | ParC5_sh110 | 49.0478993955982 | 0.557182984877337 | 1.13599765075228 |
| H3.3: K36ME2 | ParC5_sh680 | 44.9219760359209 | 0.226297413751792 | 0.503756588024617 |
| H3.3: K36ME2 | ParC5_shNT | 42.4022149802836 | 0.870600727598564 | 2.05319634364238 |
| H3.3: K36ME3 | ParC5_sh110 | 6.36217607730873 | 0.253168043970998 | 3.97926811353028 |
| H3.3: K36ME3 | ParC5_sh680 | 5.9430267791498 | 0.056615691151598 | 0.952640687237443 |
| H3.3: K36ME3 | ParC5_shNT | 5.78635204286079 | 0.232775042128554 | 4.02282889814408 |
| H3.3: K36AC | ParC5_sh110 | 2.64737952426611 | 0.214989498717091 | 8.12084163779611 |
| H3.3: K36AC | ParC5_sh680 | 3.36664614122479 | 0.103572491306058 | 3.07642938881537 |
| H3.3: K36AC | ParC5_shNT | 4.26835680568386 | 0.39084130580252 | 9.15671588846708 |

| Mark | Sample | Total Average Percent | Total Standard Deviation | Total Percent CV |
|---|---|---|---|---|
| H3: K122UN | ParC5_sh110 | 99.9900050127737 | 0.00626188221414727 | 0.00626250815103702 |
| H3: K122UN | ParC5_sh680 | 99.9873387833558 | 0.00209207248284304 | 0.00209233739821395 |
| H3: K122UN | ParC5_shNT | 99.9643539163924 | 0.00248879288237831 | 0.00248968035591955 |
| H3: K122AC | ParC5_sh110 | 0.00999498722631131 | 0.00626188221414797 | 62.6502272825709 |
| H3: K122AC | ParC5_sh680 | 0.0126612166442131 | 0.00209207248284496 | 16.5234711768489 |
| H3: K122AC | ParC5_shNT | 0.0356460836075954 | 0.0024887928823776 | 6.98195322037368 |

| Mark | Sample | Total Average Percent | Total Standard Deviation | Total Percent CV |
|---|---|---|---|---|
| H3: K9UN | ParC5_sh110 | 54.2806128642935 | 0.622484205631099 | 1.14678919928072 |
| H3: K9UN | ParC5_sh680 | 56.7455343346715 | 0.748691439463605 | 1.31938389203986 |
| H3: K9UN | ParC5_shNT | 50.7732453559818 | 1.4500319614408 | 2.85589772974787 |
| H3: K9ME1 | ParC5_sh110 | 20.8258299247851 | 0.49117102344227 | 2.35847034771815 |
| H3: K9ME1 | ParC5_sh680 | 20.2414565070795 | 0.808289697806001 | 3.9932388142291 |
| H3: K9ME1 | ParC5_shNT | 18.830539026875 | 1.07476472021067 | 5.70756216100222 |
| H3: K9ME2 | ParC5_sh110 | 19.7217421799878 | 0.806749207323725 | 4.09065892841027 |
| H3: K9ME2 | ParC5_sh680 | 18.4738348684709 | 0.472273378547257 | 2.55644473337412 |
| H3: K9ME2 | ParC5_shNT | 24.8632237795141 | 0.521741776779401 | 2.09844781757259 |
| H3: K9ME3 | ParC5_sh110 | 1.4851517473881 | 0.0487220829415379 | 3.28061311089753 |
| H3: K9ME3 | ParC5_sh680 | 1.3712196622333 | 0.00700811881118118 | 0.511086516931 |
| H3: K9ME3 | ParC5_shNT | 1.65275407730555 | 0.00799360815949236 | 0.48365381572824 |
| H3: K9AC | ParC5_sh110 | 3.68666328354553 | 0.0885233205467866 | 2.40117726351326 |
| H3: K9AC | ParC5_sh680 | 3.16795462754471 | 0.0879663678930403 | 2.77675592725322 |
| H3: K9AC | ParC5_shNT | 3.88023776032359 | 0.0573561285837802 | 1.47816015735585 |
| H3: K14UN | ParC5_sh110 | 44.5120442203437 | 0.320278630934536 | 0.719532514276567 |
| H3: K14UN | ParC5_sh680 | 44.1044100168058 | 0.58916735135194 | 1.33584680336374 |
| H3: K14UN | ParC5_shNT | 40.0717544005161 | 1.13395698494405 | 2.82981616829197 |
| H3: K14AC | ParC5_sh110 | 55.4879557796563 | 0.320278630934536 | 0.577203875028967 |
| H3: K14AC | ParC5_sh680 | 55.8955899831942 | 0.589167351351944 | 1.05404979449915 |
| H3: K14AC | ParC5_shNT | 59.9282455994839 | 1.13395698494405 | 1.89219119231786 |

| Mark | Sample | Total Average Percent | Total Standard Deviation | Total Percent CV |
|---|---|---|---|---|
| H3: K18UN | ParC5_sh110 | 95.2167306866818 | 0.0142351077099844 | 0.014950216844586 |
| H3: K18UN | ParC5_sh680 | 95.9885099597902 | 0.0835894220775481 | 0.0870827374157219 |
| H3: K18UN | ParC5_shNT | 93.925612038424 | 0.195301548218894 | 0.207932153946463 |
| H3: K18ME1 | ParC5_sh110 | 0.106814127696887 | 0.00340220208052297 | 3.18516113353245 |
| H3: K18ME1 | ParC5_sh680 | 0.0797737462254351 | 0.00170904668094688 | 2.14236733488388 |
| H3: K18ME1 | ParC5_shNT | 0.0937102889647924 | 0.00231624023760827 | 2.47170322831733 |
| H3: K18AC | ParC5_sh110 | 4.6764551856213 | 0.0122708035032649 | 0.26239540455758 |
| H3: K18AC | ParC5_sh680 | 3.93171629398441 | 0.0852332624989684 | 2.16783857546834 |
| H3: K18AC | ParC5_shNT | 5.98067767261122 | 0.193129728553747 | 3.22922817656925 |
| H3: K23UN | ParC5_sh110 | 70.0027631179303 | 0.503677355782112 | 0.71951067836221 |
| H3: K23UN | ParC5_sh680 | 72.1683880600978 | 0.45499506323969 | 0.630463109222831 |
| H3: K23UN | ParC5_shNT | 58.0170789717581 | 0.419878676228088 | 0.72371564316859 |
| H3: K23ME1 | ParC5_sh110 | 0.0352743582258045 | 0.00199245296596599 | 5.64844568740712 |
| H3: K23ME1 | ParC5_sh680 | 0.0226955577820068 | 0.000311932305049472 | 1.3744200871625 |
| H3: K23ME1 | ParC5_shNT | 0.0498695522603033 | 0.00258652550411217 | 5.18658256767841 |
| H3: K23AC | ParC5_sh110 | 29.9619625238439 | 0.503222839105318 | 1.67953897781179 |
| H3: K23AC | ParC5_sh680 | 27.8089163821202 | 0.45484620899276 | 1.63561284712699 |
| H3: K23AC | ParC5_shNT | 41.9330514759816 | 0.421398080263699 | 1.00493063450216 |
| H3: Q19UN | ParC5_sh110 | 99.9789068421028 | 0.000353041476546807 | 0.00035311595985378 |
| H3: Q19UN | ParC5_sh680 | 99.9780778010598 | 0.00171787442273466 | 0.00171825110115935 |
| H3: Q19UN | ParC5_shNT | 99.9817369934554 | 0.000813354539259927 | 0.000813503109386034 |
| H3: Q19ME1 | ParC5_sh110 | 0.0210931578972406 | 0.00035304147653675 | 1.67372509254735 |
| H3: Q19ME1 | ParC5_sh680 | 0.021922198940233 | 0.00171787442273676 | 7.83623224759631 |
| H3: Q19ME1 | ParC5_shNT | 0.0182630065446041 | 0.000813354539260582 | 4.45356320315668 |

| Mark | Sample | Total Average Percent | Total Standard Deviation | Total Percent CV |
|---|---|---|---|---|
| H3: K56UN | ParC5_sh110 | 99.9835895117533 | 0.00883918712725583 | 0.00884063791910248 |
| H3: K56UN | ParC5_sh680 | 99.9874559579673 | 7.59944469464057E-05 | 7.60039809177185E-05 |
| H3: K56UN | ParC5_shNT | 99.9664587641867 | 0.00419953572616661 | 0.00420094477495997 |
| H3: K56ME1 | ParC5_sh110 | 0.00267807241798109 | 0.00329658623976385 | 123.095485306145 |
| H3: K56ME1 | ParC5_sh680 | 0.00458657706629258 | 0.00176150284830585 | 38.4056088635551 |
| H3: K56ME1 | ParC5_shNT | 0.00897520822965116 | 0.00339427486431797 | 37.8183411177513 |
| H3: K56AC | ParC5_sh110 | 0.0137324158286955 | 0.00596322301512391 | 43.4244279339624 |
| H3: K56AC | ParC5_sh680 | 0.00795746496639205 | 0.00168633963756459 | 21.1919203500959 |
| H3: K56AC | ParC5_shNT | 0.0245660275836979 | 0.00261796893750643 | 10.6568672064983 |
| H3: Q55UN | ParC5_sh110 | 99.5823036522885 | 0.025703474596352 | 0.0258112874011238 |
| H3: Q55UN | ParC5_sh680 | 99.6345334287297 | 0.0185628068745153 | 0.0186308965734191 |
| H3: Q55UN | ParC5_shNT | 99.6465317488447 | 0.0401448597908337 | 0.0402872624729351 |
| H3: Q55ME1 | ParC5_sh110 | 0.417696347711516 | 0.0257034745963556 | 6.15362684811117 |
| H3: Q55ME1 | ParC5_sh680 | 0.365466571270311 | 0.0185628068745198 | 5.07920787665971 |
| H3: Q55ME1 | ParC5_shNT | 0.353468251155332 | 0.0401448597908332 | 11.3574160224058 |

| Mark | Sample | Total Average Percent | Total Standard Deviation | Total Percent CV |
|---|---|---|---|---|
| H3: K64UN | ParC5_sh110 | 99.9836504407478 | 0.00125609878438571 | 0.00125630418458276 |
| H3: K64UN | ParC5_sh680 | 99.9860468862766 | 0.00031423362721175 | 0.000314277478705761 |
| H3: K64UN | ParC5_shNT | 99.9770271556848 | 0.00157259783858158 | 0.00157295919204791 |
| H3: K64AC | ParC5_sh110 | 0.0163495592522049 | 0.00125609878438627 | 7.68276847718004 |
| H3: K64AC | ParC5_sh680 | 0.0139531137233611 | 0.000314233627209495 | 2.25206812930499 |
| H3: K64AC | ParC5_shNT | 0.0229728443151692 | 0.0015725978385687 | 6.84546422286202 |

| Mark | Sample | Total Average Percent | Total Standard Deviation | Total Percent CV |
|---|---|---|---|---|
| H3: K79UN | ParC5_sh110 | 91.2053653839736 | 0.337450382539272 | 0.369989617517138 |
| H3: K79UN | ParC5_sh680 | 93.997494774921 | 0.0099535165201244 | 0.0105891295762279 |
| H3: K79UN | ParC5_shNT | 83.4046120973997 | 0.404799729122259 | 0.485344537841066 |
| H3: K79ME1 | ParC5_sh110 | 0.72775982262604 | 0.0284149207364186 | 3.90443658099818 |
| H3: K79ME1 | ParC5_sh680 | 0.581928901529876 | 0.0361656013473685 | 6.21478006201275 |
| H3: K79ME1 | ParC5_shNT | 0.991392846605481 | 0.0958974223750575 | 9.67299922562577 |
| H3: K79ME2 | ParC5_sh110 | 5.75084762466884 | 0.332894787740239 | 5.78862125145263 |
| H3: K79ME2 | ParC5_sh680 | 3.57118663476446 | 0.0372800056838582 | 1.04391087603622 |
| H3: K79ME2 | ParC5_shNT | 11.9941382431337 | 0.311682978410644 | 2.59862753031942 |
| H3: K79ME3 | ParC5_sh110 | 2.24771262109631 | 0.0118250348173652 | 0.526091934813163 |
| H3: K79ME3 | ParC5_sh680 | 1.60686713619916 | 0.0888579338268725 | 5.52988681049603 |
| H3: K79ME3 | ParC5_shNT | 2.98033660653911 | 0.143054570072534 | 4.79994674959401 |
| H3: K79AC | ParC5_sh110 | 0.0683145476352197 | 0.00925447581395988 | 13.5468595406299 |
| H3: K79AC | ParC5_sh680 | 0.24252255258556 | 0.0223041815048291 | 9.19674531998849 |
| H3: K79AC | ParC5_shNT | 0.629520206321984 | 0.00812057887797401 | 1.28996318091504 |

| Mark | Sample | Total Average Percent | Total Standard Deviation | Total Percent CV |
|---|---|---|---|---|
| H3: R42UN | ParC5_sh110 | 99.9878014147578 | 0.00110562897687481 | 0.00110576386442239 |
| H3: R42UN | ParC5_sh680 | 99.9939294789188 | 0.000199183977195413 | 0.000199196069434801 |
| H3: R42UN | ParC5_shNT | 99.9769726085413 | 0.00106615874216684 | 0.00106640430726121 |
| H3: R42ME2 | ParC5_sh110 | 0.0121985852422086 | 0.00110562897687616 | 9.06358364452417 |
| H3: R42ME2 | ParC5_sh680 | 0.00607052108115965 | 0.000199183977191628 | 3.28116770419942 |
| H3: R42ME2 | ParC5_shNT | 0.023027391458708 | 0.00106615874216557 | 4.62995882133401 |
| H3: R49UN | ParC5_sh110 | 99.7957963438249 | 0.00371933481683423 | 0.00372694537555476 |
| H3: R49UN | ParC5_sh680 | 99.7505493020167 | 0.0104814060198948 | 0.0105076173447025 |
| H3: R49UN | ParC5_shNT | 99.8124212159743 | 0.00689406111850809 | 0.00690701721741696 |
| H3: R49ME2 | ParC5_sh110 | 0.20420365617508 | 0.0037193348168367 | 1.82138502635223 |
| H3: R49ME2 | ParC5_sh680 | 0.249450697983273 | 0.0104814060198929 | 4.20179462500271 |
| H3: R49ME2 | ParC5_shNT | 0.187578784025672 | 0.00689406111851502 | 3.6752883084963 |

| Mark | Sample | Total Average Percent | Total Standard Deviation | Total Percent CV |
|---|---|---|---|---|
| H3R2UN: K4UN | ParC5_sh110 | 93.1315196933278 | 0.299659911129804 | 0.321759928450166 |
| H3R2UN: K4UN | ParC5_sh680 | 93.3090627205909 | 0.301951046328877 | 0.323603128704715 |
| H3R2UN: K4UN | ParC5_shNT | 92.6718998104989 | 0.0873566358353383 | 0.0942644275276222 |
| H3R2UN: K4ME1 | ParC5_sh110 | 6.38432500131921 | 0.331828711116086 | 5.19755355574033 |
| H3R2UN: K4ME1 | ParC5_sh680 | 6.22954356066892 | 0.299513344605937 | 4.80795007995378 |
| H3R2UN: K4ME1 | ParC5_shNT | 6.44062308616699 | 0.100486272136795 | 1.56019488786135 |
| H3R2UN: K4ME2 | ParC5_sh110 | 0.286781440691752 | 0.0111784553799658 | 3.89790055904664 |
| H3R2UN: K4ME2 | ParC5_sh680 | 0.25987614996458 | 0.00252573183599593 | 0.971898281677704 |
| H3R2UN: K4ME2 | ParC5_shNT | 0.455906084372193 | 0.0293924143713459 | 6.44703270670774 |
| H3R2UN: K4ME3 | ParC5_sh110 | 0.031100242318589 | 0.00637000598937743 | 20.4821747821881 |
| H3R2UN: K4ME3 | ParC5_sh680 | 0.0328229927403202 | 0.00088975376944935 | 2.710763690833 |
| H3R2UN: K4ME3 | ParC5_shNT | 0.0506218090638774 | 0.00419122279492335 | 8.27948046983986 |
| H3R2UN: K4AC | ParC5_sh110 | 0.16627362234265 | 0.0208729223219898 | 12.5533575487853 |
| H3R2UN: K4AC | ParC5_sh680 | 0.16869457603528 | 0.0114566205494988 | 6.79133900968033 |
| H3R2UN: K4AC | ParC5_shNT | 0.380949209898059 | 0.0142965019785967 | 3.75286300827935 |
| H3R2UN: Q5UN | ParC5_sh110 | 99.8884236100647 | 0.0786683141714777 | 0.0787561874823211 |
| H3R2UN: Q5UN | ParC5_sh680 | 99.8807974204414 | 0.0813115924669174 | 0.0814086336582214 |
| H3R2UN: Q5UN | ParC5_shNT | 99.8865450424766 | 0.0698712459001896 | 0.0699506083331613 |
| H3R2UN: Q5ME1 | ParC5_sh110 | 0.111576389935304 | 0.0786683141714733 | 70.50623722195 |
| H3R2UN: Q5ME1 | ParC5_sh680 | 0.119202579558565 | 0.0813115924669183 | 68.2129470419464 |
| H3R2UN: Q5ME1 | ParC5_shNT | 0.113454957523439 | 0.0698712459001874 | 61.5850090867583 |

| Mark | Sample | Total Average Percent | Total Standard Deviation | Total Percent CV |
|---|---|---|---|---|
| H4: K5UN | ParC5_sh110 | 95.611498157016 | 0.185603110489786 | 0.194122165291232 |
| H4: K5UN | ParC5_sh680 | 96.2508402828228 | 0.138280213589635 | 0.14366650013996 |
| H4: K5UN | ParC5_shNT | 95.5761585750461 | 0.29084715949227 | 0.304309321308303 |
| H4: K5AC | ParC5_sh110 | 4.388501842984 | 0.185603110489787 | 4.22930460395079 |
| H4: K5AC | ParC5_sh680 | 3.74915971717724 | 0.138280213589632 | 3.68829881949505 |
| H4: K5AC | ParC5_shNT | 4.42384142495391 | 0.29084715949227 | 6.57453854136059 |
| H4: K8UN | ParC5_sh110 | 93.0928239722628 | 0.472593138155411 | 0.507657967596107 |
| H4: K8UN | ParC5_sh680 | 93.5838908287951 | 0.345838862119876 | 0.369549565696689 |
| H4: K8UN | ParC5_shNT | 92.0467786508531 | 0.658627881533593 | 0.715536047200375 |
| H4: K8AC | ParC5_sh110 | 6.90717602773717 | 0.472593138155417 | 6.84206014524059 |
| H4: K8AC | ParC5_sh680 | 6.41610917120494 | 0.345838862119872 | 5.39016486302903 |
| H4: K8AC | ParC5_shNT | 7.95322134914692 | 0.658627881533595 | 8.28127185978849 |
| H4: K12UN | ParC5_sh110 | 96.1395823749382 | 0.187470538535514 | 0.194998286766413 |
| H4: K12UN | ParC5_sh680 | 96.7100609011369 | 0.167898262706791 | 0.173609923458147 |
| H4: K12UN | ParC5_shNT | 95.7926628212678 | 0.272628397942868 | 0.284602588458727 |
| H4: K12AC | ParC5_sh110 | 3.86041762506176 | 0.187470538535515 | 4.85622429341478 |
| H4: K12AC | ParC5_sh680 | 3.28993909886308 | 0.167898262706799 | 5.10338512846089 |
| H4: K12AC | ParC5_shNT | 4.20733717873223 | 0.272628397942874 | 6.47983240613539 |
| H4: K16UN | ParC5_sh110 | 52.846568482819 | 0.414590102027508 | 0.784516599525843 |
| H4: K16UN | ParC5_sh680 | 65.6685531135422 | 1.9243781969596 | 2.93044098844742 |
| H4: K16UN | ParC5_shNT | 56.0927647341418 | 3.69749899468556 | 6.59175744360309 |
| H4: K16AC | ParC5_sh110 | 47.153431517181 | 0.414590102027508 | 0.879236332728927 |
| H4: K16AC | ParC5_sh680 | 34.3314468864578 | 1.9243781969596 | 5.6052930228195 |
| H4: K16AC | ParC5_shNT | 43.9072352658582 | 3.69749899468556 | 8.4211610507863 |

| Mark | Sample | Total Average Percent | Total Standard Deviation | Total Percent CV |
|---|---|---|---|---|
| H4: K20UN | ParC5_sh110 | 2.26083912084735 | 0.436286211796518 | 19.2975346088757 |
| H4: K20UN | ParC5_sh680 | 3.1048874593584 | 0.150553043191526 | 4.84890499775591 |
| H4: K20UN | ParC5_shNT | 2.97413096895123 | 0.534554171609626 | 17.9734576987417 |
| H4: K20ME1 | ParC5_sh110 | 0.379739198740985 | 0.0141860764149336 | 3.73574191496879 |
| H4: K20ME1 | ParC5_sh680 | 0.561947296367121 | 0.0838345861925319 | 14.9185852008731 |
| H4: K20ME1 | ParC5_shNT | 0.325484671650341 | 0.110204013176715 | 33.8584341369856 |
| H4: K20ME2 | ParC5_sh110 | 95.4988966647473 | 0.425427200619735 | 0.445478655228043 |
| H4: K20ME2 | ParC5_sh680 | 94.1579490205703 | 0.117076458835204 | 0.124340493875485 |
| H4: K20ME2 | ParC5_shNT | 94.5929464008041 | 0.694185912844345 | 0.733866466007918 |
| H4: K20ME3 | ParC5_sh110 | 1.84983492571089 | 0.162172176920837 | 8.76684587726194 |
| H4: K20ME3 | ParC5_sh680 | 2.12790582446832 | 0.125565940915154 | 5.90091626571529 |
| H4: K20ME3 | ParC5_shNT | 1.98214881956525 | 0.146489078525892 | 7.39041776681642 |
| H4: K20AC | ParC5_sh110 | 0.0106900899534541 | 0.00860378625842704 | 80.4837592189486 |
| H4: K20AC | ParC5_sh680 | 0.0473103992358392 | 0.00487258171825409 | 10.2991769187248 |
| H4: K20AC | ParC5_shNT | 0.125289139029071 | 0.00485514997750814 | 3.87515631054147 |
