## Supplementary Materials for "PKCδ regulates chromatin remodeling and DNA repair through SIRT6"

**SUPPLEMENTARY MATERIALS AND METHODS**

**TCGA analysis**

Patient clinical data were obtained from The Cancer Genome Atlas (TCGA) Broad Firehose for 522 Lung Adenocarcinoma (LUAD) patients [Firehose Broad GDAC – Lung Adenocarcinoma, accessed on 08/21/2021] and 528 Head-neck squamous cell carcinoma (HNSC) patients [Firehose Broad GDAC - Head-neck squamous cell, accessed on 08/24/2021]. Data was used for patients that have both PKCδ RNAseq and PKCδ CNV data (514 patients for LUAD and 516 patients for HNSCC). Patients were split into 3 groups of equivalent patient number based on PKCδ RNA-seq expression data (low PKCδ expression, middle PKCδ expression, and high PKCδ expression). Total genome wide CNV events with neutral seg. mean values between 0.3 to 0.3 were removed from the data set and remaining CNV events were counted for each patient. Box and whisker plots are shown for patient CNV counts between expression groups for each cancer type using Rstudio (v 1.4.1717) running R (v 4.1.2) and package ggplot2 (v 3.3.5).

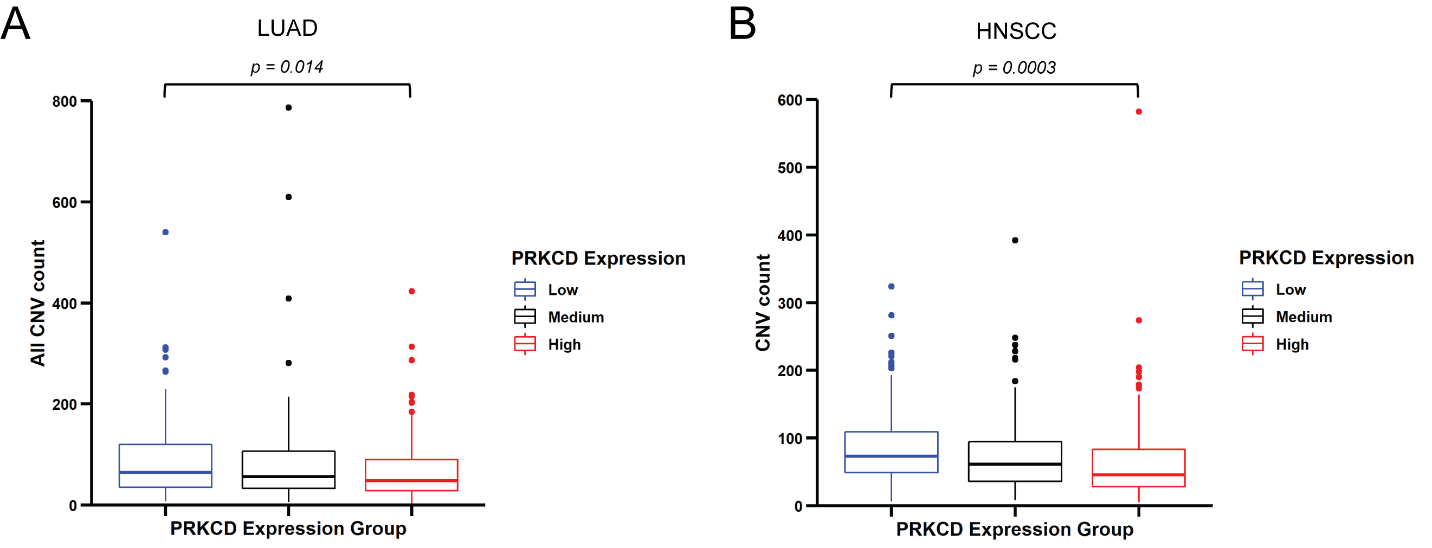

**Fig. S1. Correlation of copy number variants (CNV) and PKCδ expression in LUAD and HNSCC.** Boxplots of all CNV (gains and losses) by PRKCD (PKCδ) expression level in cancer patients with (**A**) lung adenocarcinoma (LUAD) and (**B**) head-neck squamous cell carcinoma (HNSCC). Data was generated using the TCGA database; tumors were subdivided by relative PKCδ expression level (high, medium and low). Statistics represent paired t-test.

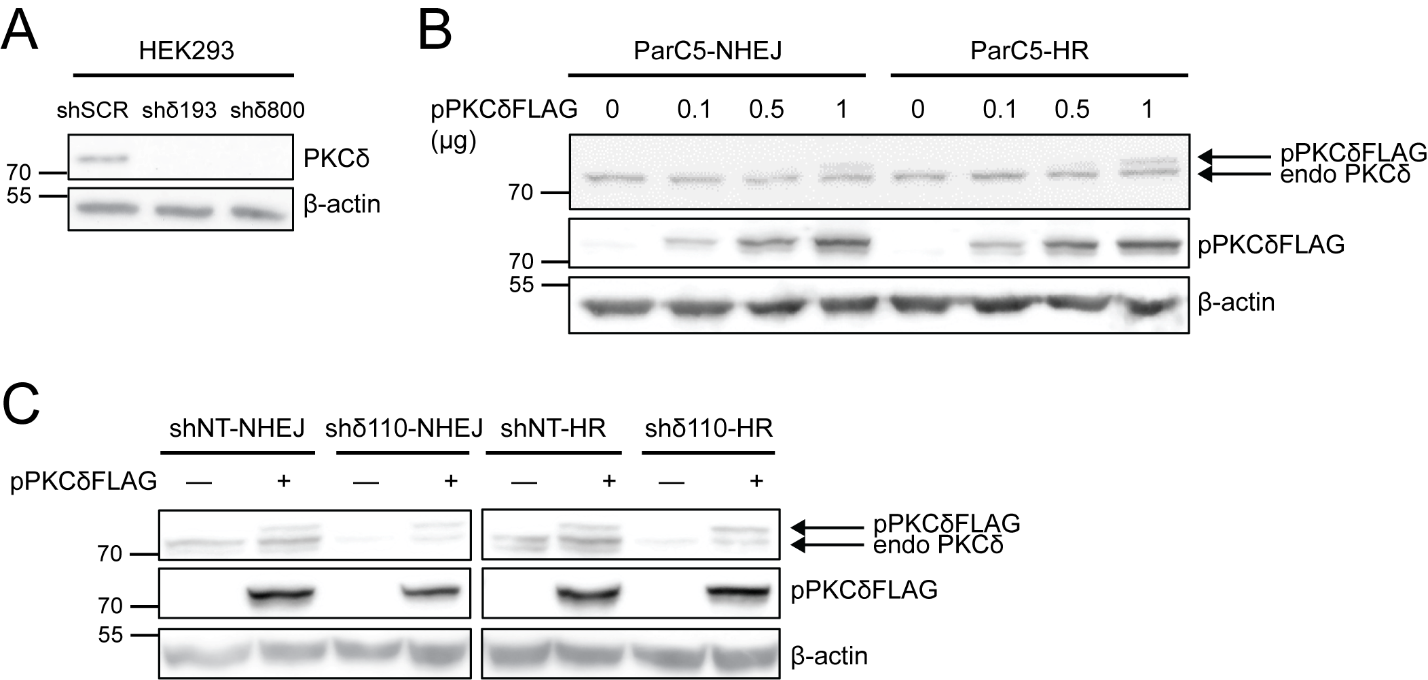

**Fig. S2. Overexpression and knockdown of PKCδ in ParC5 and HEK293 cells.** (**A**) Immunoblot analysis of HEK shSCR, shδ193, and shδ800 cells with the indicated antibodies. (**B**) ParC5 cells containing NHEJ or HR reporter were transiently transfected with empty vector (pEV, 0µg) or increasing amount of pPKCδFLAG for 72 hrs. Cells were then collected, lysed, and immunoblotted with the indicated antibodies. (**C**) ParC5 shNT and shδ110 containing NHEJ or HR reporter cells were transiently transfected with 1µg pEV (‒) or pPKCδFLAG (+) for 72 hrs. Cells were then collected, lysed, and immunoblotted with antibodies against PKCδ (top blots), FLAG-tag (middle blots), and β-actin control (bottom blots).

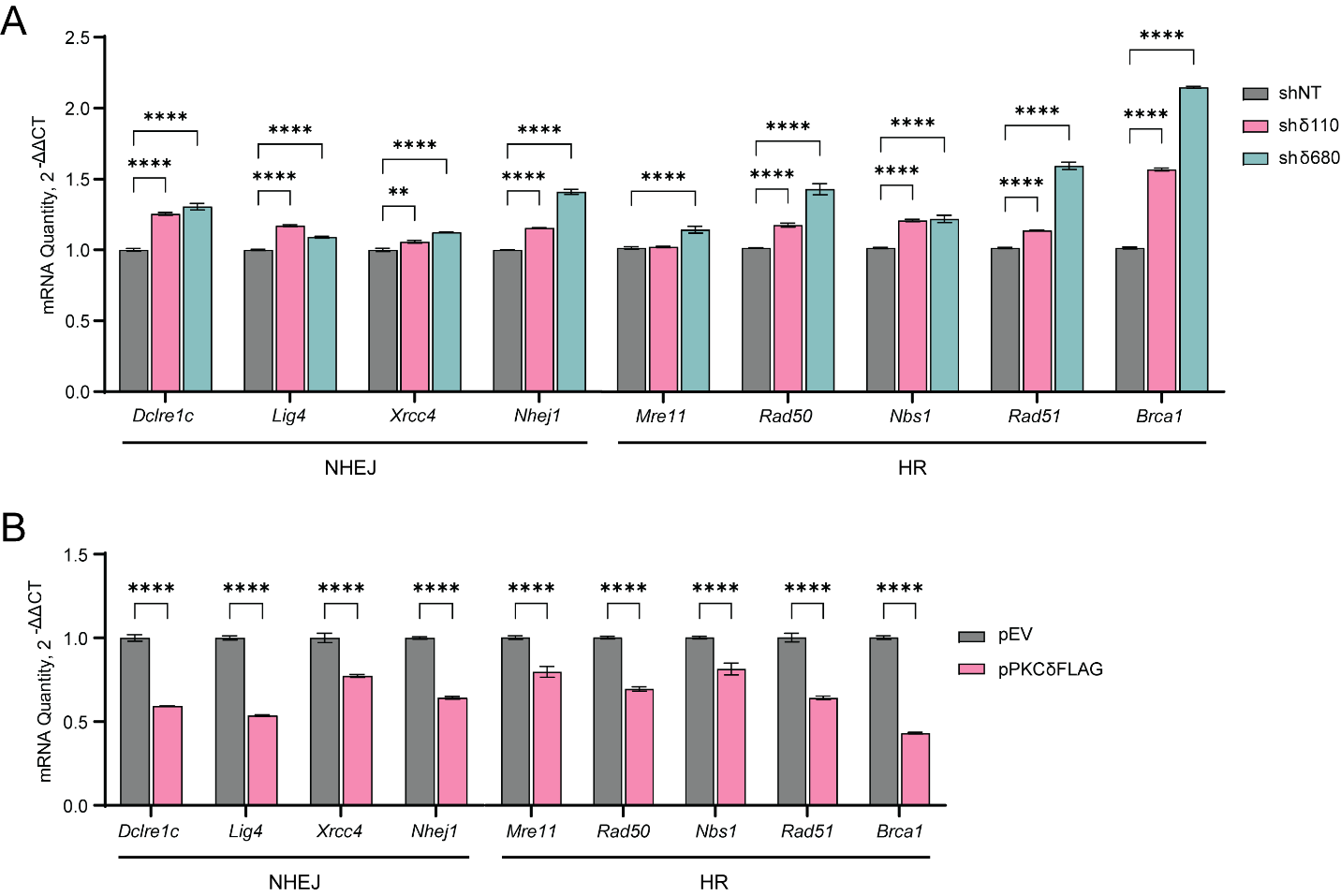

**Fig. S3. PKCδ regulates expression of genes required for DNA repair**. (**A**) The relative expression of NHEJ and HR genes in ParC5 shNT, shδ110, and shδ680 cells was assayed by qRT-PCR. (**B**) The relative expression of NHEJ and HR genes in ParC5 cells transiently transfected with empty vector (pEV) or pPKCδFLAG for 48 hrs was assayed by qRT-PCR. Shown is data (mean ± SEM) from at least three independent biological replicates. *Dclre1c-*Artemis, *Nhej1*-XLF. Statistics represent two-way ANOVA followed by (A) Dunnett’s or (B) Sidak’s multiple comparisons within each fraction to their corresponding shNT or pEV control. ***P* < 0.01 and *****P* < 0.0001.
